# Study of Impacts of Two Types of Cellular Aging on the Yeast Bud Morphogenesis

**DOI:** 10.1101/2024.02.29.582376

**Authors:** Kevin Tsai, Zhen Zhou, Jiadong Yang, Zhiliang Xu, Shixin Xu, Roya Zandi, Nan Hao, Weitao Chen, Mark Alber

## Abstract

Understanding the mechanisms of cellular aging processes is crucial for attempting to extend organismal lifespan and for studying age-related degenerative diseases. Yeast cells divide through budding, providing a classical biological model for studying cellular aging. With their powerful genetics, relatively short lifespan and well-established signaling pathways also found in animals, yeast cells offer valuable insights into the aging process. Recent experiments suggested the existence of two aging modes in yeast characterized by nucleolar and mitochondrial declines, respectively. In this study, by analyzing experimental data it was shown that cells evolving into those two aging modes behave differently when they are young. While buds grow linearly in both modes, cells that consistently generate spherical buds throughout their lifespan demonstrate greater efficacy in controlling bud size and growth rate at young ages. A three-dimensional chemical-mechanical model was developed and used to suggest and test hypothesized mechanisms of bud morphogenesis during aging. Experimentally calibrated simulations showed that tubular bud shape in one aging mode could be generated by locally inserting new materials at the bud tip guided by the polarized Cdc42 signal during the early stage of budding. Furthermore, the aspect ratio of the tubular bud could be stabilized during the late stage, as observed in experiments, through a reduction on the new cell surface material insertion or an expansion of the polarization site. Thus model simulations suggest the maintenance of new cell surface material insertion or chemical signal polarization could be weakened due to cellular aging in yeast and other cell types.

**Significance Statement:** Aging yeast exhibits two modes with different bud shapes. Experimental data analysis reveals that control of growth rate and bud size is more robust in cells aging in a spherical budding mode than in cells aging in a tubular budding mode. A computational model was developed and used in combination with experiments to test the hypothesized mechanisms underlying different types of budding in aging cells. Model simulations suggest that localized growth is sufficient to generate tubular budding and its aspect ratio can be stabilized through the regulation of chemical signals with an expanding polarization site or a decline on the new cell surface material insertion. Proposed mechanisms of morphological changes in aging yeast can be present in other cell types.

## Introduction

Cells can reach their replication limit and stop further division due to the cellular aging process, which is related to many diseases including cancer, osteoarthritis and atherosclerosis. Understanding the fundamental mechanisms underlying cellular aging can benefit the treatments of those age-related diseases. Cellular aging impacts many processes controlling biophysical properties and biochemical signaling dynamics in individual cells. *Saccharomyces cerevisiae*, also known as yeast cells, undergo a cellular aging process after multiple cell cycles. Yeast cells can reproduce through a process called budding. In each cell cycle, a projection is first formed at a specific location on the cell membrane determined by some signaling molecules. Later it separates from the mother cell and becomes a daughter cell after reaching a certain size. The yeast budding process has been extensively studied, including the governing gene regulatory network and structural changes at subcellular level, to understand asymmetric growth and cell division (1–5). Recently, it has also been used to study cellular aging by tracking their cell cycles until they stop proliferation permanently (6–15). Cellular changes that accompany the aging process have been identified mostly in the mother cell (5, 10, 16–24).

In fact, due to the cellular changes inside mother cells as they age, bud growth and overall shape also become different. When yeast cells are young, they always generate symmetric and rounded bud shapes. The final bud size is also quite robust at the moment when the bud separates from the mother cell. Different bud shapes have been observed in some mutations. For example, cells with defective actin polarization or disrupted septin polymerization or stability tend to produce elongated buds with an aspect ratio, the long axis to short axis, greater than 1.5 (25–28). However, a recent study shows that the elongated bud shape can also be observed as a natural outcome when yeast cells get old (6, 15). In fact, for aging yeast cells, the bud development in one cell cycle has been identified to differentiate into two distinct trajectories due to either chromatin instability, called Mode 1, or mitochondrial decline, called Mode 2. As a result, the aged yeast cells would either produce an excessive elongated bud with a very high aspect ratio and large size, or a relatively small and spherical bud in those two different modes, respectively (6, 7, 15, 29). However, how different bud shapes are generated in those two modes due to aging remains unclear.

The budding process involves biochemical signaling dynamics and mechanical changes. Within each cell cycle, it begins with the polarization of a signaling molecule Cdc42 and several growth-associated proteins on the cell surface. This polarization is controlled by the interaction involving multiple proteins such as Cdc42, Cdc24, Bud3, and Rga1, as well as mechanical feedbacks provided by the cell wall integrity pathway (30, 31) and the cell geometry (32), and in the end a single cluster of activated Cdc42 is usually achieved serving as the spatial cue for the bud formation. As the biochemical polarization stabilizes, septins are recruited to form a “septin cloud” within the Cdc42 cluster (33). Consequently, the septin cloud assembles into a ring structure, called the septin ring, which determines the exact bud site. Within the bud site, actin cables are constructed, and their integrity is maintained by Bni1, an effector of Cdc42, to promote the secretion and delivery of new cell surface materials toward the bud site. During the bud formation, secreted cell wall modifying enzymes alter the crosslinking in the network of beta-3-glucan, the main building blocks of the yeast cell wall, allowing cell wall remodeling to accommodate new materials and new crosslinking (34).

The entire budding process, starting from the polarization of Cdc42 and growth-associated protein in G1 phase to the abscission of the daughter cell in M phase, usually takes approximately 90 minutes for younger cells. As the cell ages, the budding process also becomes much longer, and it can take more than 300 minutes for an old cell that is nearly at the end of its lifespan (6). However, what causes the slow-down of the budding process for an aged yeast cell and how the two different bud shapes are formed for older cells remain mysterious. It could be related to the production and delivery rate of cell surface materials, or the cell wall extensibility that determines the cell wall deformation. Furthermore, changes in the mechanical properties may also play a role since it has been suggested that aged yeast cells have the cell wall elasticity altered where the bud scars are formed, and the cell surface usually becomes stiffer in comparison to ones with less bud scars (35). Therefore, based on the recent experimental observations in (6, 15) and major components identified to be involved in the yeast budding process, we hypothesize that the delivery rate and location of new cell surface materials, under the regulation of polarized chemical signals, determine the bud growth rate and the final bud shape. Cellular aging may affect the size of the polarization site of chemical signals and maintenance of the polarization site, giving rise to different modes of budding in old cells.

Previously, we developed a 3D coarse-grained particle-based model to study the effect of changes of mechanical properties along the cell surface on the bud formation process (36). The sphere, representing the combined cell wall and membrane of a mother yeast cell in the model, is discretized into a triangulated mesh in 3D. Vertices in each triangle are connected by linear springs to capture the in-plane elasticity, whereas neighboring triangles sharing a common edge are connected with each other through bending springs to represent the out-of-plane elasticity of the cell surface (37). Moreover, the model assumes that turgor pressure remains constant throughout one cell cycle before the bud separates. Using this computational framework, it has been shown that a higher stretching to bending stiffness ratio in the bud region than that in the mother cell is sufficient for bud emergence (38).

In this study, a combined experimental analysis and modeling simulation approach is used to test hypothesized mechanisms regulating cell cycle length and giving rise to different bud shapes due to cellular aging in yeast. By measuring bud growth rate, size and shape from experimental images obtained at different ages of yeast cells fixed in a microfluidic channel, it is found that budding always follows a linear growth regardless of cell age, and cell cycle length becomes much longer in old cells. Cells in different aging modes exhibit different robustness in maintaining the bud size even when cells are young. For old cells that give rise to tubular budding, the bud aspect ratio only increases at the early stage of each cell cycle and remains constant while the bud keeps growing at the late stage. By applying the calibrated model developed in (36) and coupling it with the chemical signaling model of Cdc42 pathway, we have identified the linear bud growth can be achieved by a constant rate of delivery of new cell surface materials. Moreover, tubular bud shape can only be obtained by a spatially localized insertion of new cell surface materials, but not the nonhomogeneous mechanical properties over the cell surface. Simulation results also predict the stabilized aspect ratio of tubular budding can be resulting from the diminishing polarization of Cdc42, suggesting cellular aging may affect the maintenance of the polarizing signals.

## Results

### Experiments suggest two modes of cellular aging with different growth rates and final sizes and shapes

Bud growth of wild type yeast has been shown to produce a linear increase in the bud surface area when the mother cell is relatively young (39). However, the budding process of old mother cells has not been well studied. The age of a yeast cell can be determined via the number of replications the cell has. Hence, it is called the replicative age (9, 40). The assay is usually analyzed by tracking the number of daughter cells a yeast cell has produced. Alternatively, since each yeast cell has a different number of total replications, the approximated age is usually determined by the lifetime percentage (current number of replications divided by total number of replications), and it has been found that the cell cycle time is quite robust among yeast cells at a similar lifetime percentage (6).

Budding of individual yeast cells was tracked using experimental images throughout the lifetime of each cell (14) (see Materials and Methods for more detail) (Fig. 1). It has been shown that aging yeast cells can develop into two different bud shapes, with one more tubular, denoted as “Mode 1” aging, which is driven by nucleolar decline, and the other more spherical, denoted as “Mode 2”, which is driven by mitochondrial deterioration (6). We measure the bud surface area and calculate the bud aspect ratio for both young and old mother cells to investigate the bud shape formation and budding growth rate as cells age in different modes (Fig. 2).

**Figure 1.**
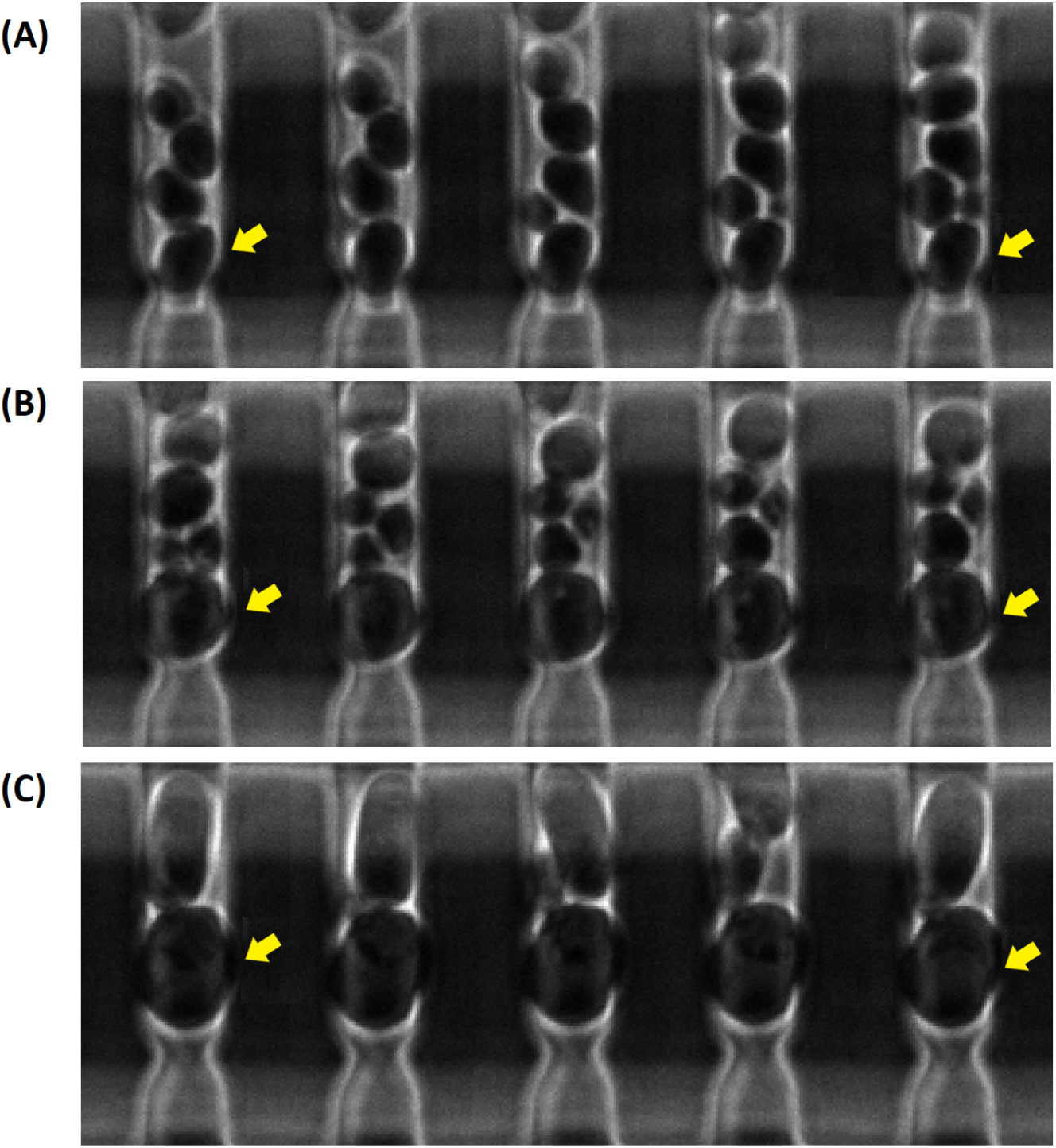
Sample experimental images depicting the budding process. The progression is oriented from left to right where the leftmost image represents the beginning of the budding process. (A) A young yeast cell producing spherical buds. (B) An aged yeast cell in mode 2 gave rise to a spherical bud. (C) An aged yeast cell in mode 1 gave rise to a tubular bud. Yellow arrows indicate the mother cells tracked. Time between successive snapshots is (A) 15 minutes, (B) 30 minutes, (C) 30 minutes except the rightmost image was captured after 180 minutes since the previous snapshot.

**Figure 2.**
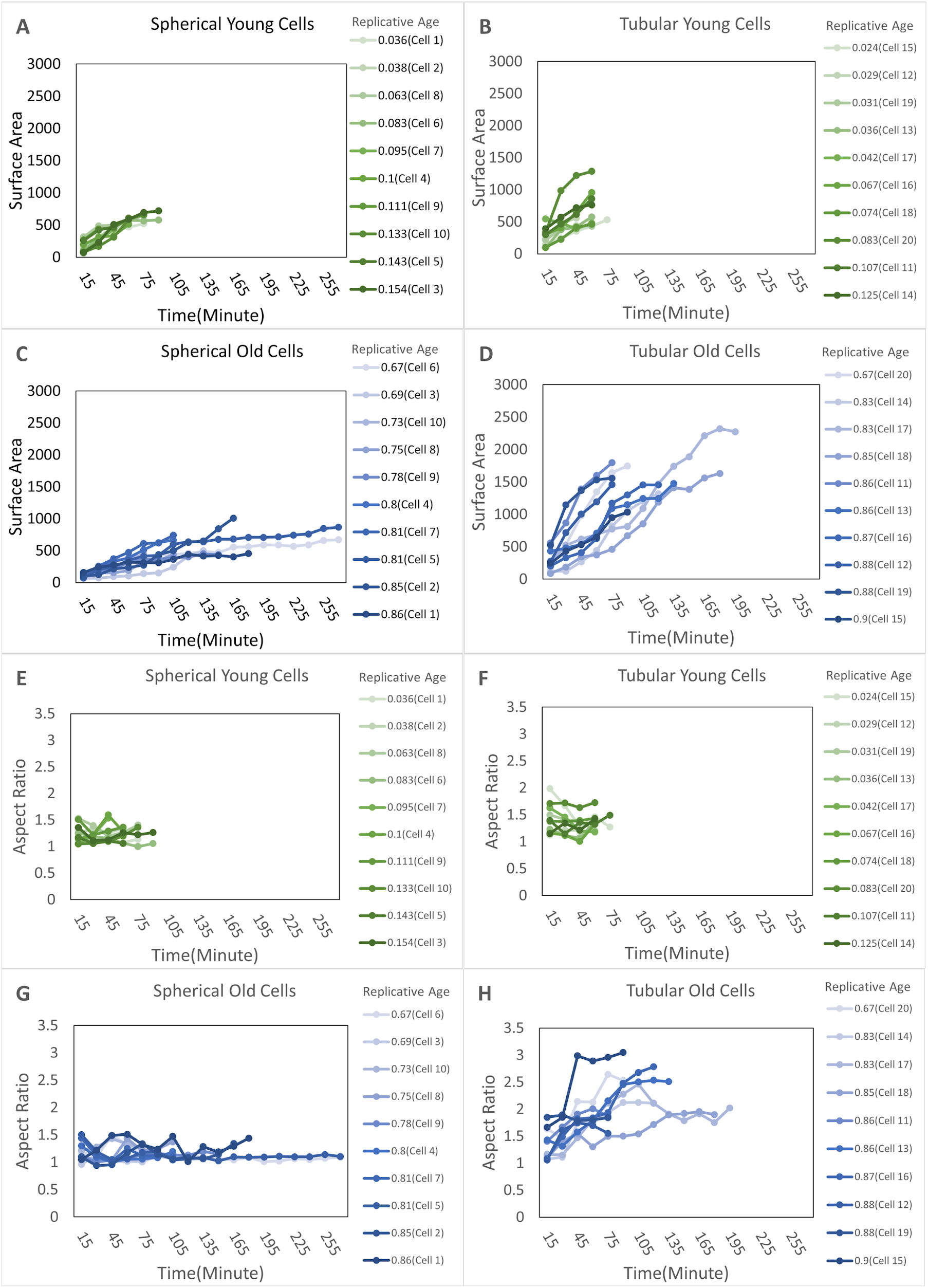
Image analysis of bud growth rate and aspect ratio of cells at different ages in experiments. (A) Time history of surface area for spherical buds from young yeast cells which still give rise to spherical buds when they are old. (B) Time history of surface area for spherical buds from young yeast cells which give rise to tubular buds when they are old. (C) Time history of surface area for spherical buds by the same mother cells as (A) when they are old. (D) Time history of surface area for tubular buds of the same mother cells as (B) when they are old. (E) Time history of aspect ratio for spherical buds from young yeast cells which still give rise to spherical buds when they are old. (F) Time history of aspect ratio for spherical buds from young yeast cells which give rise to tubular buds when they are old. (G) Time history of aspect ratio for spherical buds by the same mother cells as (A) when they are old. (H) Time history of aspect ratio for tubular buds of the same mother cells as (B) when they are old.

### Spherical budding reaches similar final bud size at different ages with growth rate reduced in older cells

We first focus on cells that always give rise to spherical budding regardless of cell age, i.e. Mode 2. Budding process is initiated fast on young mother cells and the bud surface area undergoes a linear growth (Fig. 2A), which is consistent with the observation in previous literature (39). The growth rate is quite consistent among different samples, and they all reach similar final sizes within about 75-100 minutes after bud initiation. This suggests young mother cells in Mode 2 experience very similar cell cycles and buds generated have similar shapes. As mother cells age, the cell cycle becomes significantly longer. Some of the buds grow much more slowly after bud initiation and some old mother cells even take a much longer time to initiate the budding process (Fig. 2C). However, the final bud size obtained remains similar to those obtained by young mother cells. The aspect ratio of buds, which is quantified as the ratio between the longest axis and shortest axis, stays close to 1 no matter how old mother cells are, and it also remains constant during budding for both young and old mother cells (Fig. 2EG). In summary, this data indicates, in Mode 2, older mother cells have much longer cell cycles than young cells, while maintaining the spherical bud shape throughout the budding process and reaching a robust final bud size.

### Tubular budding of aging cells shows large variation in both bud size and growth rate

In contrast, cells in Mode 1 have very different behavior even when they are young. Although young mother cells in this aging mode also generate spherical buds, the growth rate varies a lot from sample to sample (Fig. 2B). Some of the buds grow linearly at rates similar to cells in Mode 2. Some grow much faster with short cell cycles, and some grow much more slowly. Although most samples still grow linearly, the variation among young cells in Mode 1 is much larger than that in Mode 2. Similar behavior is also observed on the bud aspect ratio (Fig. 2F). The variation among samples is large, while it remains constant within individual cell cycles and the overall average is still close to 1, indicating spherical shape is still maintained in individual cell cycles for young mother cells in Mode 1.

As mother cells in Mode 1 get older, bud initiation becomes faster on average (Fig. 2D) and bud shape becomes asymmetric right after the initiation (Fig. 2H). For most of the samples we collect, buds still grow linearly, and the bud aspect ratio increases rapidly only during the early stage followed by a saturation, while the bud keeps growing. Large variation is observed on the aspect ratio from sample to sample (Fig. 2H). All tubular buds collected reach much larger final sizes than those generated by young cells and the cell cycle of aging cells becomes much longer. In comparison to cells in Mode 2, which always generate spherical buds at similar size at different ages, the final size of tubular budding shows a large variation among samples (Fig. 2A-D), indicating a less effective size controlling mechanism governing aged cells in this mode. The growth rate of tubular budding also varies significantly from sample to sample, but the average rate is similar to that in young cells (Fig. 2B, D).

Based on the comparison between young and old mother cells in those two modes, we notice that yeast budding behavior starts to diverge in terms of the shape’s robustness when they are still young, although the average bud size and shape remain similar for the two aging modes. As mother cells age, the divergent behavior becomes more visible and buds with different sizes and shapes are generated, while the bud growth remains linear in both modes. Such linear growth on the bud surface area, observed in both spherical and tubular budding, suggests a model that new cell surface materials are delivered and inserted into the bud region periodically at a roughly constant rate throughout the budding process. Moreover, this cell surface material insertion rate changes as cells get older as shown in spherical budding or tubular budding. Next, we develop a computational framework based on the model developed in (36), coupled with polarizing chemical signal Cdc42, to investigate the mechanism underlying the growth rate change and bud shape formation in different modes due to cellular aging. We assume the cell surface material insertion rate to be constant and the growth event is introduced periodically in the model as described in detail in Materials and Methods. Furthermore, we calibrate the increase in the bud surface area per growth event in the model using the temporal bud growth rate obtained from experiments. Such calibration on the bud growth trajectory is implemented for both young and older mother cells, as well as spherical budding and tubular budding (See Supplementary Information). We also assume the new cell surface materials are only added to the polarization region of Cdc42 within the bud surface. We apply this coupled model to test whether nonhomogeneous mechanical properties or nonhomogeneous cell surface growth can give rise to the tubular bud shape as observed in Mode 1 in experiments.

### Model simulations predict the mechanism underlying different bud shape formation due to cellular aging

Recently, it has been noticed that wild type yeast cells could undergo distinct aging-driven differentiation processes (6, 15). One cell life cycle will keep progressing toward either nucleolar decline with more tubular bud shape, or mitochondrial decline with more spherical bud shape (6, 7). Our new experimental data suggests those two aging trajectories diverge at a young age. The mechanism underlying different bud shape formation due to cellular aging remains unclear. During the budding process, some chemical signaling is polarized on the cell membrane, which rearranges the actin cables to follow a similar polarized distribution inside the cell (34, 41–44). Consequently, new materials are delivered along the polarized actin cables to the cell surface, leading to bud growth. Such new material insertion in the bud surface may also change the mechanical properties locally or the local mechanical properties may affect the new material insertion. Extensive studies have been conducted to understand the dynamics of those critical signaling networks and the mechanism to establish cell polarization (2, 30, 34, 45, 46). By coupling the mechanical model we developed previously in (36) with the chemical signaling model describing one of the major signaling pathways in yeast, Cdc42, we study the mechanism underlying different bud shape formation due to cellular aging.

### Growth restricted to the bud tip leads to tubular budding

In our chemical submodel, we utilize a spatial signaling gradient *u* to represent the gradient of some extracellular molecular stimuli and to determine a preferred polarization site of intracellular signaling molecules including Cdc42. This allows us to control the local concentration of Cdc42 and its associated growth proteins in a simplified way (see Materials and Methods). The gradient *u* is described by the following formula: *u*_*T*_ = *u*_*min*_ + (*u*_*max*_ − *u*_*min*_) · ((*H*_*T*_ − *H*_*min*_) / *H*_*total*_)^*n*^ where larger *n* results in a sharper gradient and smaller *n* results in a shallower gradient. The growth region within the bud surface is chosen to be the area with concentration of Cdc42 greater than some threshold value. Varying the slope of the transition from its maximum to the minimum associated with the gradient *u*, fixed throughout the simulation, will lead to a change in the growth region. We ran the simulations with *n* = 2 and *n* = 8 respectively. With *n* = 2, due to the almost linear spatial gradient *u*, the distribution of the signaling molecule was less polarized and its concentration over the entire bud was higher than the threshold value at the early stage (Fig. 3A). Therefore, new materials were inserted almost evenly over the entire bud surface, and a spherical budding was generated (Fig. 3A). With *n* = 8, the spatial gradient *u* was decaying rapidly from its maximum at the bud tip to the rest of the bud surface, and therefore the signaling molecule was more concentrated near the tip. By using the same threshold value, the growth region in this case became much smaller and new materials were only inserted to a subregion around the tip, giving rise to a tubular bud (Fig. 3B). Therefore, our coupled model can generate both spherical budding and tubular budding by employing different gradients *u* as the spatial cue. For a shallower spatial cue, the growth region is large relative to the bud size at the early stage (Fig. 3D) and a spherical bud is obtained. For a steeper spatial cue, since the growth region determined by the chemical signal is sufficiently small relative to the bud size throughout the budding process, a tubular bud is obtained with an increasing aspect ratio (Fig. 3C). Overall, this suggests growth restricted at the bud tip is sufficient to give rise to tubular budding. Cellular aging may affect the polarization of the signaling molecule through different biological processes including restricting the diffusion and activation in space or localizing the delivery and insertion of the new materials within the bud surface, such that the growth becomes more restricted to the bud tip, and it generates tubular budding.

**Figure 3.**
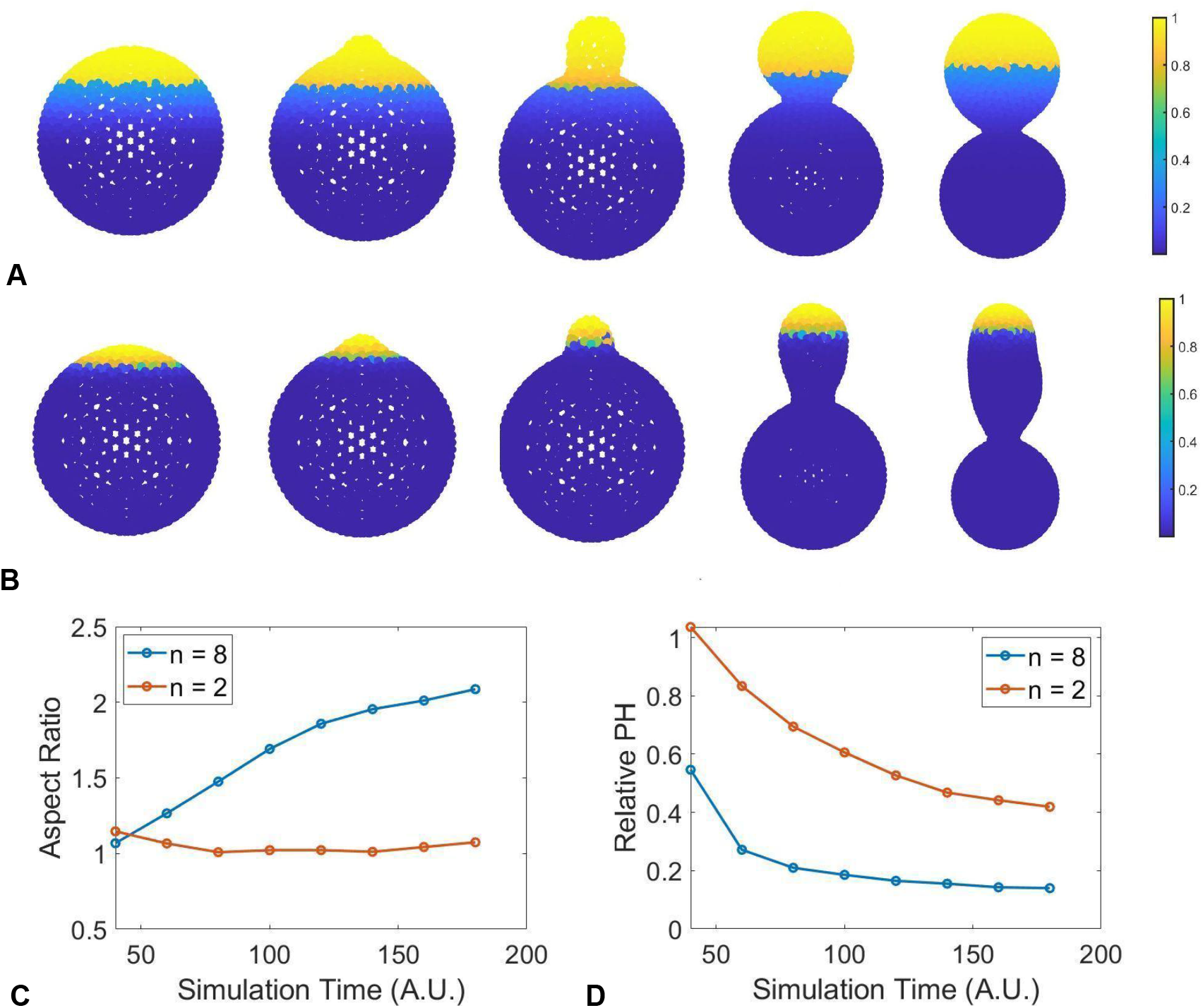
Sample simulations where the mechanical model is coupled with a chemical signaling model controlling the size of the polarization zone which determines the location of cell surface material insertion. Polarization shown here is a time-dependent process. Growth, restricted to the bud site, characterized by material insertion is introduced based on the local concentration. Such concentration must exceed a specific threshold value, yellow color represents locations where material insertion is possible while blue color represents locations whose current chemical concentration is below a threshold value, 0.8 · *Conc*_*max*_, where *Conc*_*max*_ is the maximal value of concentration over the whole cell surface. (A) depicts a case where the polarization is controlled by setting *n* = 2, where *n* is the constant determining the sharpness of the polarization. (B) depicts a case when *n* = 8. The more polarized the signal directing material insertion, the more elongated the resulting bud becomes. (C) Corresponding aspect ratio during bud formations. (D) Corresponding polarization height (PH), based on the concentration threshold described above, during bud formations.

### Kinetics of stabilizing the tubular bud shape

In simulations, the cell wall over the entire bud surface can rearrange itself locally, which is modeled by a stochastic re-meshing approach, called edge swapping (ES) (36, 47). The goal of edge swapping is to energetically relax the system after addition of new nodes. To see the effect of edge swapping on the bud shape, we first test the scenario of spatially biased growth with the growth region fixed at the tip of the bud and then let the entire bud surface relax. To test the effect of the relaxation time, i.e., the number of edge swapping steps, we run the simulations with 25, 75 or 125 edge swapping steps following each growth event. The simulation results show that the bud shape changes from being tubular to spherical as the number of steps increases (Fig. 4A). This suggests that the tubular bud shape can only be obtained when the growth time significantly surpasses the relaxation time. In other words, more materials must be added before the system reaches equilibrium. Thus, the tubular bud shape can only be obtained under the nonequilibrium conditions, not as a minimum energy structure, under the spatially biased growth.

**Figure 4.**
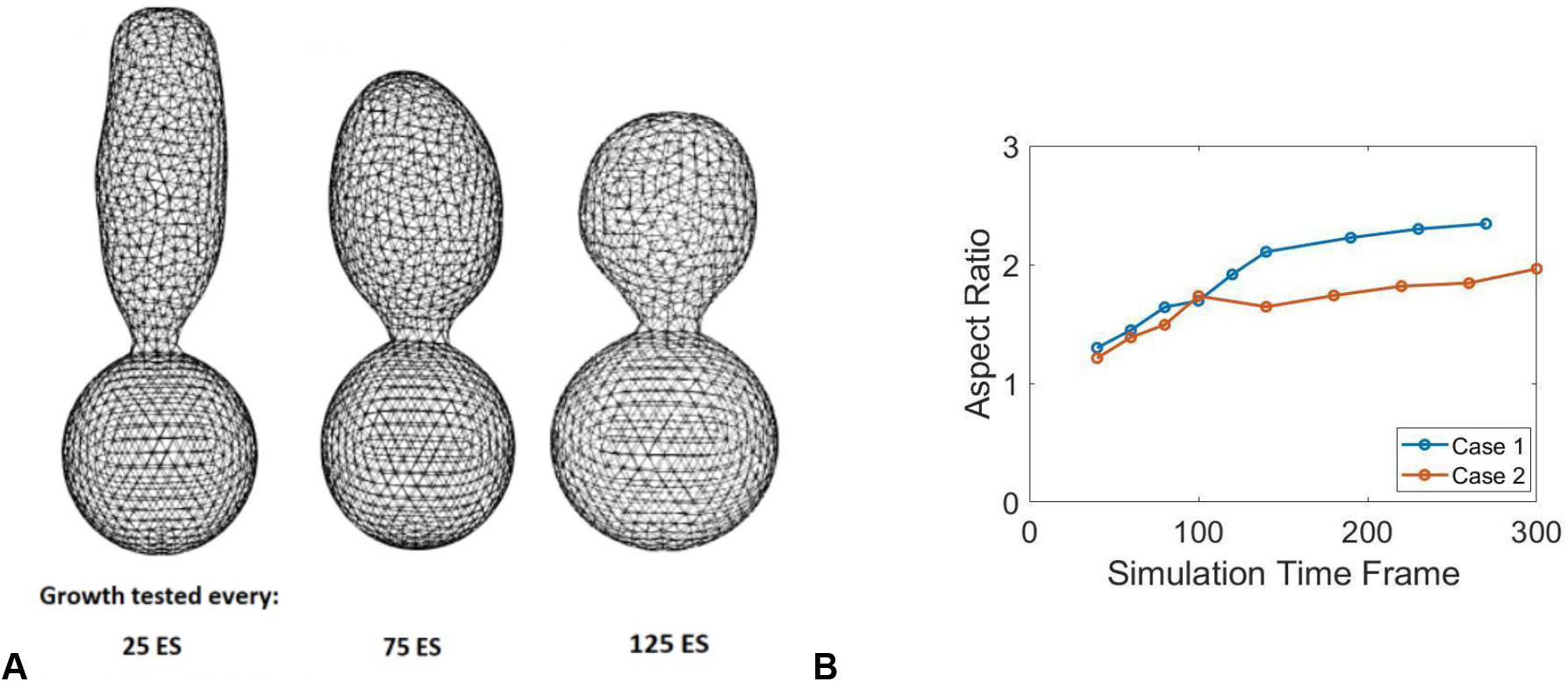
Impact of Edge Swapping (ES) on bud shape. (A) Simulations of budding with spatially biased growth and with 25, 75, or 125 ES steps between successive growth events. As the number of ES steps increases, the relaxation time extends, leading to a progressively more spherical bud shape. Note that the material insertion is confined to a small region at the bud tip. (B) Increased edge swapping steps between successive growth events can stabilize the aspect ratio of the bud shape. Case 1 and 2 represented the simulations where the edge swapping steps between successive growth events changed from 25ES to 50ES after 150 and 100 simulation time frames, respectively.

As observed in experiments, the tubular budding has its aspect ratio increasing only at the early stage and then it is maintained while the bud keeps growing until the end of the cell cycle (Fig. 2H). This suggests that the cell wall rearrangement may be enhanced at the later stage of budding, which can stabilize the aspect ratio. So, we test the case with an increasing number of edge swapping steps between successive growth events during budding. By Implementing edge swapping more frequently at the later stage of budding, the aspect ratio can be almost stabilized as reaching a steady state (Fig. 4B). Therefore, the stabilized aspect ratio of tubular budding in the late stage may result from a balance between the new cell surface material insertion and the cell wall rearrangement to relax the energy.

Another hypothesis is that the spatially restricted growth might be dynamically changing during budding such that the growth is more localized at the beginning and then spreads into a larger region, under the guidance of polarized biochemical signaling molecules. In the following section, we study a time-varying growth region to understand this alternative mechanism for maintaining the aspect ratio in tubular budding.

### Reduction in polarizing signal stabilizes the aspect ratio of tubular buds

Via experimental observations, distribution of the polarizing signaling molecules, especially Cdc42 and its associated growth proteins, switches from being polarized during bud emergence to a more spreading distribution within the bud, followed by localization near the bud neck at the end of one cell cycle (48). We hypothesize that the spatially restricted growth region changes over time in a similar manner, determined by the biased spatial gradient *u*, which is assumed to be also changing over time in the chemical signaling submodel. In particular, the parameter *n* in the formula of *u* is assumed to be a function of time, *n*(*t*). This temporally changing spatial cue is used to shrink at the early stage and then expand the growth region determined by the polarized signal (Fig. 5A-B). The aspect ratio of the bud first increases in simulations and then stays constant while the bud keeps growing (Fig. 5C). The stabilization of the aspect ratio starts exactly at the time when the signal becomes less polarized. It was also shown that the size of the growth region, relative to the bud size, decreases first and then stays the same (Fig. 5D). For *n*(*t*) being reduced over a longer time period in the function of *u*, representing the gradient changing from being steep to shallow, the aspect ratio is maintained at a higher level (case 2 in Fig. 5), compared to the case with more rapidly decreasing *n*(*t*) (case 1 in Fig. 5). This suggests that cellular aging in yeast may affect how long the signaling polarization can be maintained after it is established and how fast the polarized distribution vanishes.

**Figure 5.**
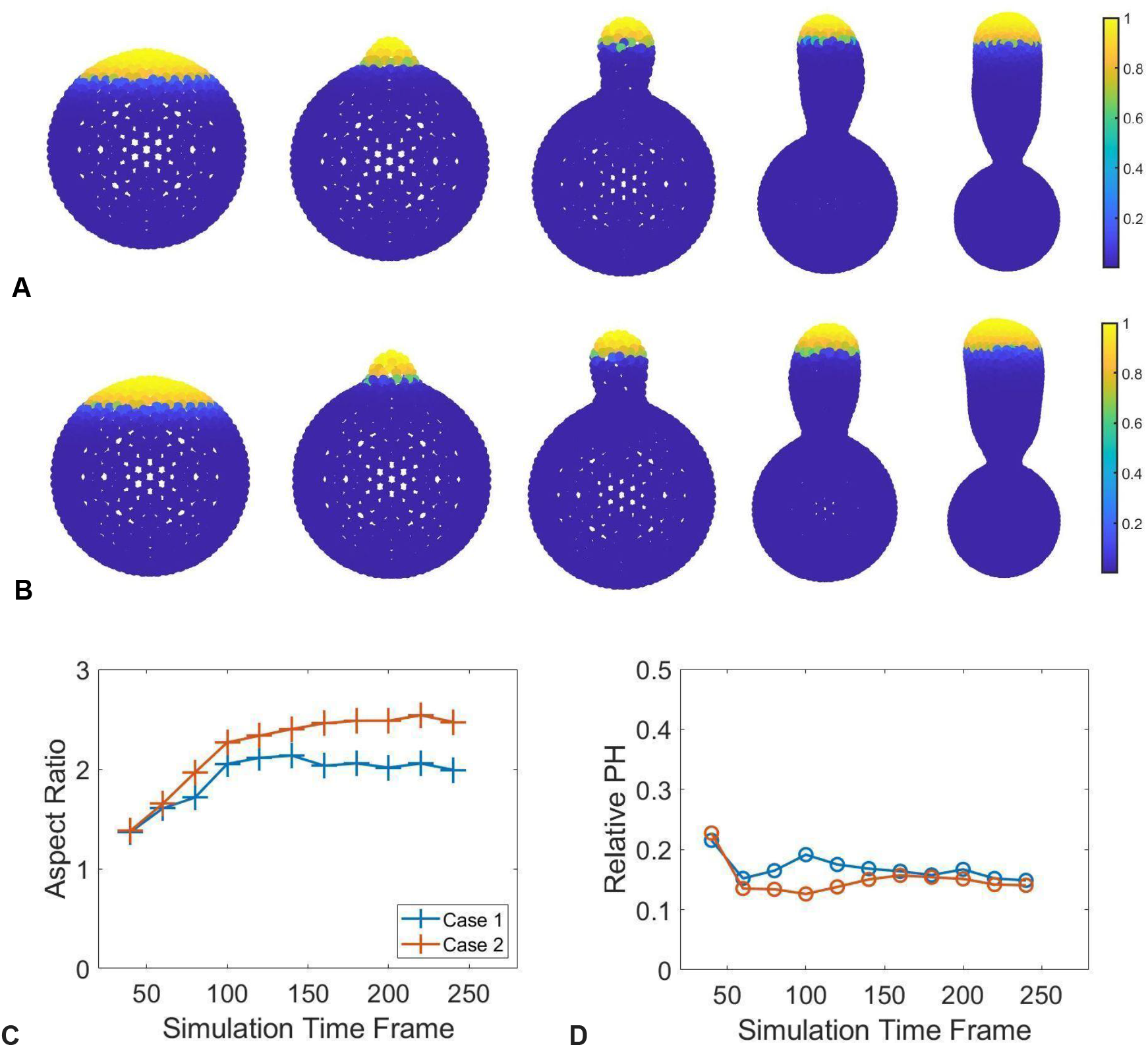
(A, B) Sample simulation snapshots demonstrating the change of the polarization pattern during bud development corresponding to Case 1 and 2 in (C, D), respectively. From left to right, each image corresponds to simulation time frame 2, 20, 40, 80, and 160, respectively. (C) Aspect ratio of the bud produced. Case 1 represents the scenario where the scaling factor *n* starts with 5 and increase to 12 in 5×10^3^ simulation time steps, and then decrease to 7 in 5×10^3^ simulation time steps. Case 2 is of the identical setup to Case 1 except the latter decrease of the scaling power *n* completes in 1×10^4^ simulation time steps. (D) Corresponding relative PH (relative height of the region eligible for material insertion to the height of the bud). The maintenance of the aspect ratio is directly linked to the relative PH in terms of stabilization of such values.

## Discussion

In this study, a combination of experimental data analysis and biologically calibrated multiscale computational mechano-chemical model was used to test hypothesized mechanisms underlying different modes of bud shape formation and changes in bud growth rates of aging yeast cells. Experimental data analysis has shown that cells aging in different modes have different levels of robustness in controlling bud size and growth rate even at the young age. This data was also used to calibrate a multiscale model coupling particle-based Monte Carlo type submodel and signaling submodel, to study the bud emergence without aging effects. The coupled model included a novel chemical signaling submodel representing Cdc42 signal in the form of a system of reaction-diffusion equations solved on the triangular mesh on the growing 3D cellular surface. Cdc42 signal can achieve polarized distribution at the early stage of one cell cycle and is responsible for regulating the rate and location of new cell surface materials. The simulated polarized chemical signal was used in the mechanical submodel to guide the spatially biased growth. The model was calibrated to reproduce the linear growth of budding by applying a constant insertion rate of new cell surface materials within the Cdc42 polarization site. The model was first calibrated to achieve experimentally observed linear cellular growth rate and then used to test hypothesized mechanisms underlying the tubular bud shape formation and maintenance of its aspect ratio while the size keeps increasing at the later stage of one cell cycle.

Analysis of experimental images suggests that the budding dynamics is quite different in two aging modes even when cells are young, although the bud surface area always increases linearly in both modes. For spherical budding, average bud size remains similar at different ages, while the variation among different samples becomes larger and the growth rate is reduced in aging cells (Fig. 2C). For tubular budding, even when cells are young and still produce spherical buds, the variation on bud size among different samples is much larger, indicating less robust bud sizes obtained in this mode (Fig. 2B). Variation of bud sizes and growth rates among different samples increases significantly in aging cells with tubular budding (Fig. 2D). We calibrated the mechano-chemical model to achieve a specific experimentally observed linear growth rate of budding. Model simulation results indicate that non-uniform mechanical properties of the cell surface are not sufficient to generate asymmetric bud shapes. They further predict that the spatially biased growth which may occur in aging cells is required to give rise to tubular budding. Such tubular bud shape can only be maintained under insufficient cell wall modification, computationally represented by employing a limited number of Monte Carlo cell surface remeshing on the bud surface as a nonequilibrium (Fig. 4). By coupling with the dynamic polarizing chemical signal Cdc42, the model can generate more biologically relevant bud shapes with aspect ratio maintained at the later stage of tubular budding. Such maintenance relies on an expanding polarization site of Cdc42 in the late stage of one cell cycle, suggesting cellular aging may affect the maintenance of chemical signal polarization.

One possible extension of the studies described in this paper would be to consider the bud initiation guided by the polarizing biochemical signals in individual cell cycles. In our current model, the bud location was given manually as one initial condition and fixed throughout the simulation. However, it was suggested that structural components required to initial budding were delivered by actin cables whose distribution was dependent on the polarization of the signaling molecules (49, 50). However, it was still under debate how early and how quickly the biochemical signals became polarized and how this process was changing during aging. The existence of the septin ring would also affect the chemical signaling distribution, since it would reduce the diffusion as a physical barrier (51–53). It is worth looking at the polarizing signals throughout one cell cycle in old cells experimentally.

Another interesting question to study was how the linear growth in bud surface area was achieved mechanistically. In our current model, the new material insertion rate into the cell surface is assumed to be constant, which might be an outcome of the mechanical properties and the regulations of governing biochemical processes. This growth rate is maintained within one cell cycle and reduced as mother cells get older. It would be interesting to link the new material insertion rate to polarizing signals in the model to understand the mechanism.

It would also be interesting to simulate the late stage of the budding process including the cell division, and to explore the size control mechanism as mother cells get old. Our experimental data showed that the final bud size remained similar for cells at different ages in the mode of spherical budding, whereas the final bud size varied a lot among different samples in the mode of tubular budding even when mother cells are still young. This suggested that the bud size control may not be affected by aging in spherical budding, but it becomes less robust even in young cells with tubular budding. It is known that aging cells may undergo ribosomal DNA instability, leading to dysregulation in ribosomal biogenesis involved in cell size control (10, 16). It has been observed that the polarized signaling molecules lost the polarization distribution near the tip and became concentrated near the septin ring before the cell division occurred. It was then followed by the shrink of the septin ring, leading to the complete separation of the bud from the mother cell. Both biochemical signals and mechanical components were rearranged when the final bud size was achieved. Our mechano-chemical model could be applied to understand the underlying mechanism. By simulating the entire cell cycle of yeast budding, it would be also possible to extend the model to study the budding process with multiple generations and understand how cellular aging affected the critical biological processes dynamically to give rise to different bud shapes.

Overall, the experimental data analyzed here suggests that cells can age in two different modes with different levels of robustness in controlling bud size and growth rate from the start. This indicates that aging mechanisms may take effect in cells already at young ages and have an impact on the size control. Model simulations predict potential mechanisms for bud morphogenesis during cellular aging which can be verified in future experiments on the localization and maintenance of Cdc42 polarization in aging cells. This mechano-chemical model can be also adapted to study aging of other cell biology systems in the context of cell growth and cell division.

## Materials and Methods

Experimental procedures and techniques can be found in SI. A detailed description of the mechanical model and mechano-chemical coupled model is also provided in SI, including the modeling calibration, numerical method and additional modeling analysis.

### Mechano-chemical Computational Model

In this section we describe coupling of different submodels in space leading to a multi-scale model of cell budding. The novelty of the chemical signaling submodel is in its implementation on a growing 2-dimensional cell surface in 3-dimensional space. Namely, a quasi-steady state solution of a system of reaction-diffusion partial differential equations representing cell signaling is numerically calculated on the deforming triangular mesh generated by the mechanical model determining cell surface. Mechanical properties and the insertion region of new cell surface materials are updated based on the spatial gradient of the chemical signal to follow a similar polarized pattern within the bud surface. This is a two-way coupling in the sense that the chemical signal impacts cellular mechanical properties and spatially biased growth, whereas the resulting bud shape provides a deforming domain for the signaling submodel. This multi-scale model is applied to study the interplay between the cellular physical properties and the biochemical cues affecting bud shape formation and maintenance.

The signaling submodel involves Cdc42 similar to (38). It is known that at the beginning of a budding cycle, Cdc42 initially was distributed almost uniformly along the cell surface and cytosol. Upon activation, multiple clusters of Cdc42 form on the cell surface and, eventually, they evolve into a single polarization site due to the self-promoting activation and competition between clusters on the acquisition of polarization-promoting microstructures (2, 27, 36). In the chemical signaling model we introduced, the competition between Cdc42 clusters is neglected since it only happened over a short time period and it was assumed that the location of the single polarization site was determined by a spatial cue *u*. In addition, we assume that the inactive Cdc42 and other growth-associated proteins were well-mixed in the cytosol at the beginning. To establish a polarization on the cell surface *Γ*, we utilized the following reaction-diffusion partial differential equations studied in (54):

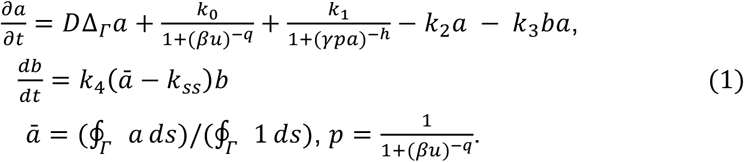

The novelty of the current paper is that we solve this system of equations on a growing surface in 3-dimensional space. Here, *a* represents the concentration of activated Cdc42 associated with cell surface *Γ*, and *b* represents the concentration of its negative regulators, such as the protein kinase Cla4p which is activated by Cdc42 and inhibits it (55). ∮_*Γ*_ * *ds* denotes the integral over the surface of cell membrane *Γ*, and Δ_*Γ*_ denotes the Laplace-Beltrami operator on *Γ*. An initial biased spatial cue *u*, for example the pheromone gradient in extracellular space, was introduced to determine the location of the polarization site. More details of each individual term can be found in the Supplemental Information in (38). In particular, *u* is varied in time to represent different levels of polarization at different stages of one cell cycle. The maximal value of *u* was set to be achieved at the tip of the bud, whereas its minimum was set to be achieved at the point on the surface of the mother cell directly opposite to the bud site. The gradient *u*(*t*) is modified to be *u*(*t*) = *u*_*min*_ + (*u*_*max*_ − *u*_*min*_) · ((*H*_*T*_ − *H*_*min*_)/*H*_*total*_)^*n*(*t*)^, where *n*(*t*) is the scaling factor such that higher values give rise to a steeper transition between the maximum and minimum of *u*. In particular, *n*(*t*) is non-static, and increases at the early stage followed by a reduction to generate a time-varying polarization, instead of a fixed constant.

The numerical scheme for solving Eqs. (1) is described in detail at the end of the Supplementary Information file for the present paper. By simulating the signaling reaction-diffusion submodel over the bud surface provided by the mechanical model, a spatially based distribution of Cdc42 is established and maintained on the cell surface (Fig. S1A). The level of polarization including the size of polarization site is determined by the sharpness of the initial cue *u* and self-enhancement. At the same time, solution of the reaction-diffusion equation is used to determine the location for cell surface materials insertion, leading to the growth of the bud surface. Namely, when the solution on the modeled cell surface patch exceeds a threshold value, such patch becomes eligible for cell surface expansion by adding new triangles into the existing cell surface mesh. (For detailed description of the cell surface expansion method please see (36)). The robustness of the reaction-diffusion signaling submodel is tested on a cell surface mesh that undergoes Monte Carlo remeshing periodically. The results showed that the numerical scheme is stable. Furthermore, the stability is maintained when applied to a dynamical cell surface mesh where both the Monte Carlo remeshing and cell surface expansion are introduced (Fig. 3 & 5).

## Data, Materials, and Software Availability

Codes used for model simulations and data analysis are available at Github (https://github.com/librastar1985/BuddingCode_UCR).

## Acknowledgments

The authors acknowledge partial support from the NSF Grant NSF Grant DMS-2029814 (to MA and WC) and the NSF Grant DMS-1853701 (to WC). M.A. and K.T. were also partially supported by the DOE Office of Biological and Environmental Research, through project 74860. ZZ and NH were supported by National Institutes of Health Grants R01AG056440, R01AG068112, and R01GM144595. RZ was supported by the NSF DMR-2131963 and the University of California Multicampus Research Programs and Initiatives (Grant No. M21PR3267). Model simulations were performed using the computer clusters and data storage resources of the UC Riverside HPCC, which were partially funded by NSF grant MRI-2215705.

## Supplementary Information for

**This PDF file includes:**

**Supplementary text**

**Figures S1 to S9**

**Tables S1 to S5**

**Legends for Movies S1 to S2**

**SI References**

**Other supplementary materials for this manuscript include the following: Movies S1 to S2**

### Imaging

Time-lapse images were taken by using a Nikon Ti-E inverted fluorescence microscope equipped with an EMCCD camera (Andor iXon X3 DU897) and a CFI plan Apochromat Lambda DM 60X oil immersion objective (NA 1.40 WD 0.13MM). The microfluidic device loaded with yeast cells was fitted onto the motorized stage upon objective. Images were acquired every 15 min for a total of 90 hours or more with the exposure setting of 50 ms for phase channel. Detailed method has been described previously (1–4).

### Image analysis

Image analysis was performed using the image processing package Fiji (https://imagej.net/software/fiji/). Samples were selected from time-lapse data according to the cell cycle time and cell geometry for each individual bud. The aspect ratio extracted from the experimental images was calculated using the long and short axes of the cells of interest via manual measurement. The measurement was performed using the “straight” line function in Fiji to measure the longest and shortest axes, approximately mutually orthogonal, of each bud. In addition, the width of the mother-bud junction is measured to approximate the width of the septin and chitin ring structure. Such width was then used to calculate the approximate initial bud surface area. We identified the initiation of a bud by one video frame (15 minutes per frame) before the first visual observation of a visible protrusion on the cell surface. Due to the 2D nature of the image, we assumed that the mother cells and buds attain axial symmetry along the axis parallel to the mother–bud orientation, and the shape is equivalent to that of an ellipsoid. Thus, the bud surface and bud volume were approximated using the ellipsoidal surface area and volume formula. Young cell data was collected at the first possible generation in each data set. Old cell data was collected at the fourth generation before cell death for the same mother cell.

### Model setup

The mother cell is represented in the computational model as a coarse grained 2D triangulated surface, embedded in 3D space, with dynamical re-meshing and growth capability. Re-meshing, local membrane deformation and bud growth are carried out following a Markov chain process. Chemical signaling submodel developed in this paper is based on computational implementation of a system of reaction-diffusion equations on the growing cell membrane in 3-dimensional space and cell polarization is obtained in the model due to a biased spatial gradient of some extracellular chemical substance of interest. For detailed description of the chemical signaling model, please see (5). The reaction-diffusion equation describing a chemical signaling network is numerically solved on the triangular mesh on the growing cell surface obtained in the mechanical model with re-meshing and mechanical relaxation. That is to say, a typical simulation has the reaction-diffusion equation initiated and solved for every 25 re-meshing-then-mechanically-relaxed operation. When numerically solving the reaction-diffusion equation, a total number of 5000 iterative steps are involved each time to reach the quasi-steady state.

### Mechanical submodel

The mechanical properties of the model surface are represented by appropriate choices of energy potential describing an elastic material. Movement of each node of the triangulated surface is governed by the force, calculated via the negative gradient of the corresponding energy potentials, acting on it. For detailed model descriptions regarding the choice of energy potential, re-meshing algorithm and growth algorithm, please refer to (6).

### Coupled mechano-chemical model

The mechano-chemical model coupling is established via mutual feedback performed by allowing a cell membrane to evolve for a given time period and afterward the chemical concentration to be updated on the new 2-dimensional surface in a 3-dimensional space. Spatial information such as the global x,y,z-coordinates and local surface curvature are utilized in the chemical signaling model, a system of reaction diffusion equations, to determine the change in local chemical concentration. A conversion from global coordinates into curvilinear coordinates enables the proper treatment of the Laplace-Beltrami operator in the reaction diffusion equations. The local chemical concentration obtained is utilized by the mechanical model to determine the local mechanical properties and/or the suitable location for new cell surface material insertion driving the growth of a cell.

### Model calibration

We have used a newly developed 3-dimensional discrete particle-based model, which takes into account anisotropicity in cell surface mechanical properties and dynamical re-meshing of the model geometry, to propose and test the necessary condition for bud emergence. We proposed that the cell surface at the budding site must undergo weakening (or softening), particularly in the resistance to bending deformation, for a bud to emerge. In this work we calibrated the model by performing a systematic study on model parameter perturbation to observe effects on the budding cycle duration and compared the results with experimental data. We first investigated how mechanical properties of the bud affected the bud growth trajectory in the computational model to identify possible changes on mechanical properties leading to different cell cycle length for spherical budding. In particular, we perturbed the cell surface material insertion period (*NΔt*), stretching stiffness (*k*_*s*_), and bending stiffness (*k*_*b*_) of the bud surface, and the threshold value on surface expansion (*γ*) required to allow new material insertion in the computational model to simulate single cell budding. For each simulation, we quantified the bud surface area throughout the budding and plotted the growth trajectory in time, as shown in Fig. S1. Notice that, there was no mechanism on the termination of the budding process incorporated in the model, therefore simulations were stopped when a similar bud-to-mother cell surface area ratio as the experimental data was reached. We focused on the budding initiation and the linear growth rate of the budding trajectory when perturbing those parameters.

First of all, within the parameter regime such that a bud could occur, by assuming homogeneous mechanical properties and new material insertion rate within the bud region, as well as allowing growth to occur everywhere within the bud, the growth trajectory always followed a linearly increasing trend and the bud shape was always spherical in simulation results. By perturbing the cell surface material insertion period, i.e., the number of time steps between successive growth events *N*, a shorter insertion period had the bud growth initiated earlier. As studied in our previous work (6), bud emergence was determined based on the visibility of a protrusion from the cell surface with a volume of at least 5.5% of the total cell volume. Simulation results showed that, once the bud started to grow, the bud growth rate was dependent on the new material insertion period, i.e., smaller *N* gave rise to a larger budding growth rate (Fig. S1A).

Changing the mechanical properties such as stretching and bending stiffness only influenced the budding initiation time without affecting the growth rate (Fig. S1B). Reducing the weakening on the bending stiffness or enhancing the weakening on the stretching stiffness within the bud surface got the budding initiated earlier without changing the bud growth rate afterwards. This was surprising since mechanical properties should affect the bud surface area growth introduced by new material insertion. It was possible that the spatially homogeneous material insertion under different mechanical properties led to similar growth rate due to the sufficient relaxation implemented between successive growth events. Perturbing the threshold value for surface expansion (*γ*) affected the bud initiation mostly, but only a marginal effect was observed on the budding growth rate after the initiation (Fig. S1C). Larger *γ* led to a longer time to start the bud growth. More specifically, as shown in Fig. S1, when *γ* = 0.025, all simulated samples started fast budding simultaneously, which was about 5 min after the cell division in the previous cell cycle. For *γ* = 0.075, the average initiation time was about 20 min. When *γ* = 0.1, the bud didn’t grow significantly until 40 min after the cell division in the previous cell cycle. The growth rate after the bud was initiated was not affected by *γ* too much, i.e., the budding growth trajectory exhibited similar growth rate for *γ* = 0.025, 0.075, 1 and the variation on the growth trajectory among different simulated samples using the same parameter set was mainly due to the initiation time. The bud size from the simulations also remained similar for different *γ* (Table S1). Therefore, the threshold value for surface expansion in the algorithm affected the bud initiation significantly without changing the growth rate.

**Figure S1.**
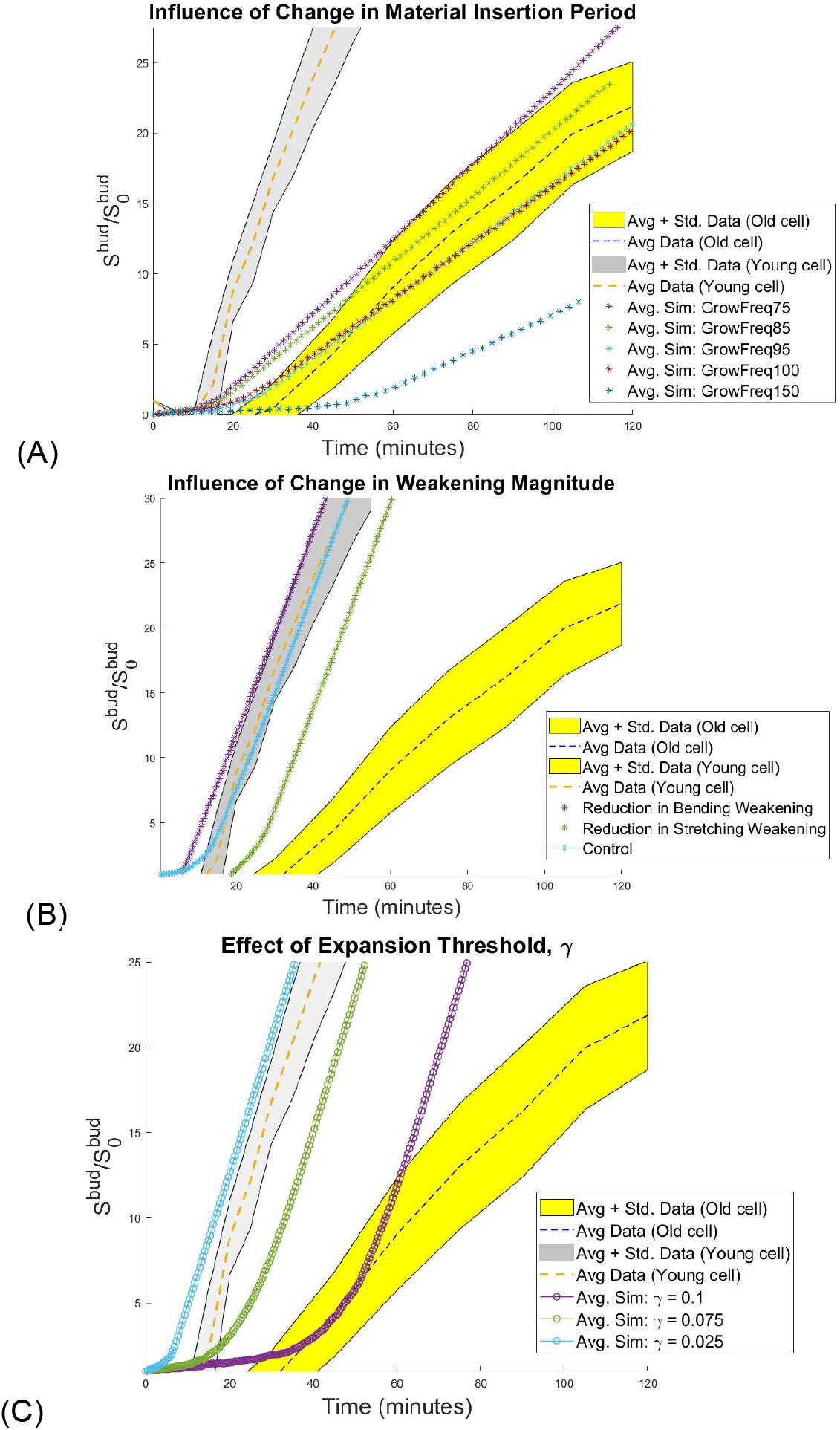
Possible changes in Mode 2 with spherical budding during cellular aging. Effect of changes in cell surface material insertion period (A) and changes in the stretching or bending stiffness (B). The y-axis represents the ratio of the bud surface area with respect to the estimated initial bud site area enclosed by the septin and chitin ring. Experimental data is collected from aged cells. (C) Effect of changes in the threshold value of surface expansion (*γ*). Increasing *γ* led to delayed bud initiation. Bud growth trajectories showed marginal differences after bud initiation. Data presented here is extracted from young cell data.

The simulation results suggested that, for spherical budding, the growth rate was only affected by the cell surface material insertion period, whereas the bud initiation time could be affected by the mechanical properties, or the new material insertion rate, or the cell surface expansion threshold within the bud region. This indicated that the new cell surface materials were delivered less frequently to the bud region as cells aged in Mode 2. The delay of the bud initiation could be due to the less frequent new cell surface material insertion, or enhanced weakening on the mechanical properties, or a larger allocated space required for new material insertion. Therefore, appropriate parameter values were chosen in the model such that similar bud initiation time and growth rate as the experimental data could be obtained (Fig. S2).

**Figure S2.**
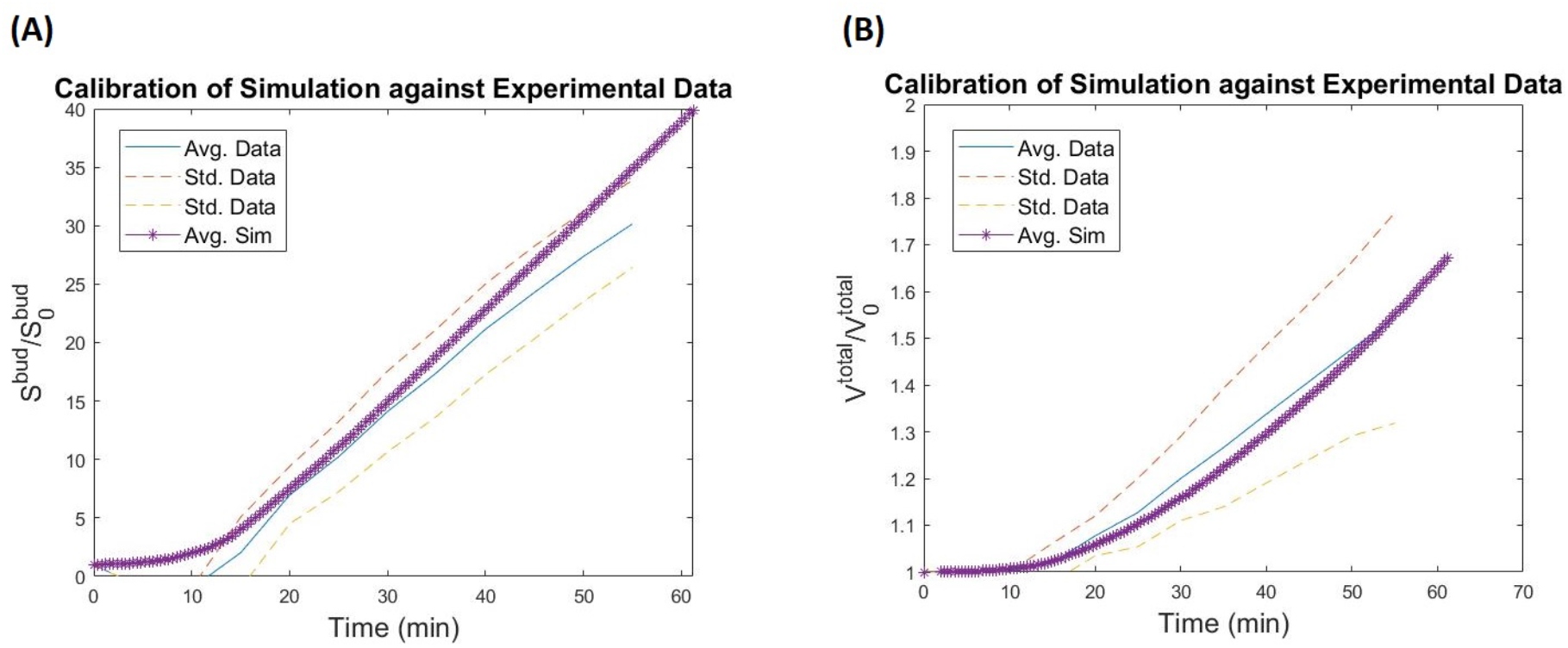
Calibration against the experimentally observed budding cycle duration for young wild type yeast cells. (A) Ratio of bud surface area with respect to initial bud surface area, and (B) Total cell volume with respect to initial total cell volume. 10 simulations were performed to ensure statistical significance. The average simulation trajectory is compared to the observed data range. Observed data range is presented as the mean trajectories with standard deviations.

More specifically, 10 simulations were performed for each parameter set during calibration for statistical significance. Our calibration results suggest that using the parameter set (α_*s*_, α_*a*_, α_*b*_) = (0.75, 0.135, 0.75) coupled with the surface area expansion threshold *γ* = 0.05 and a single cell surface expansion event accepted every 1250*Δt* unit time, where *Δt* = 8×10^−5^ *min* is the simulation time step size, nine out of ten simulations results fell close to or within the experimentally observed bud cycle duration for the young wild type yeast cell. Despite the fact that this choice of parameter set produced results in close agreement with the experimental data, this does not imply that there are no other working parameter sets due to a lack of precise time-series data on the mechanical properties of budding yeast cells.

### Stability analysis of the mechano-chemical model

In order to solve differential equations involved in the chemical signaling submodel on the cell surface obtained in the mechanical submodel, we adopted the locally discontinuous Galerkin method. This method utilized the local coordinate transformation and local curvature information, which can effectively capture the effect of surface curvature on the diffusion operator, and is ideal to handle the Laplace-Beltrami operator in the 3D setting. However, since our mechanical model incorporates stochastic components that can alter the connectivities between surface nodes in the mesh, it is essential to first investigate the stability of the numerical method on such a dynamical mesh. To evaluate the stability, we fixed a parameter set and solved the differential equations on a spherical surface with multiple trials. The results from multiple trials suggested that the method is stable without any sign of numerical “blow ups” or similar problems (Fig. S3). It is worth noting that in this validation attempt, we did not require the polarization to converge to a steady state before triggering the growth algorithm since our interest here was to verify the stability.

**Figure S3.**
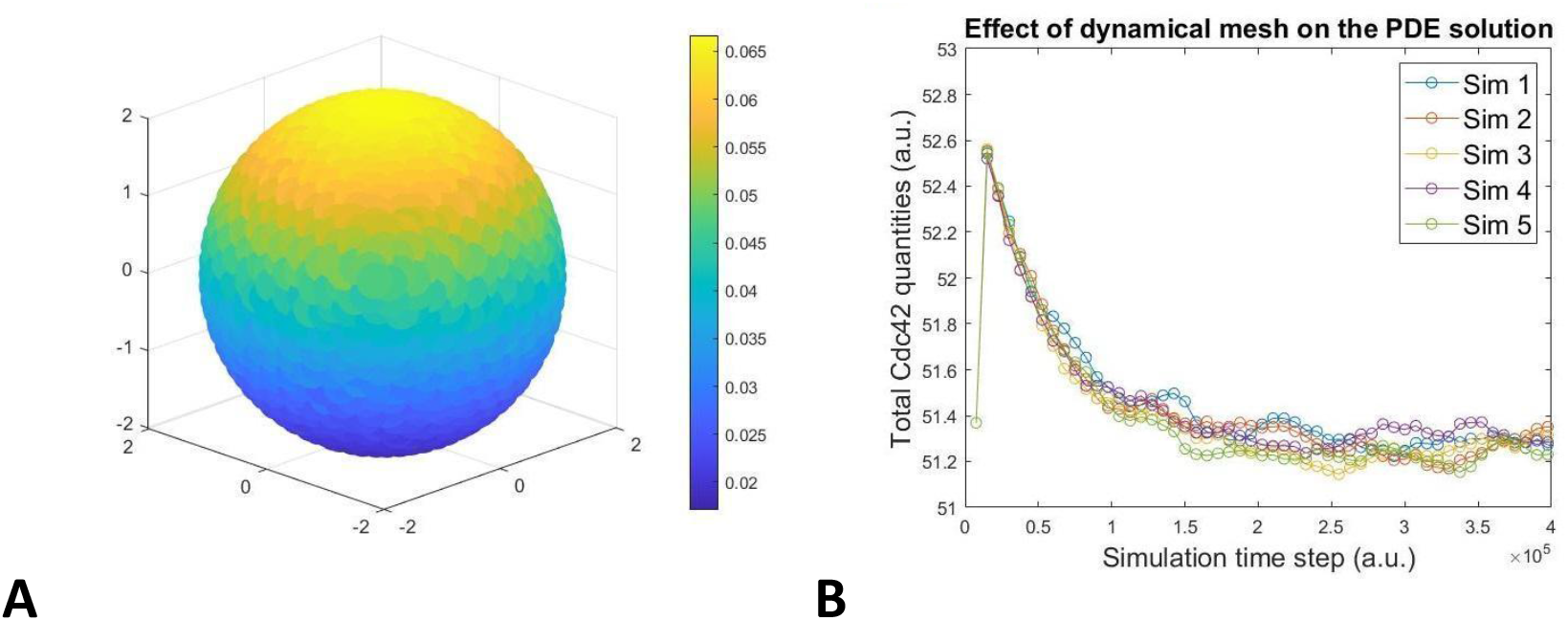
(A) A simulated polarized distribution of active Cdc42 over the triangulated sphere. The size of the polarization zone can change depending on the input parameter used for solving the differential equations. (B) The total quantity (in arbitrary units) of the Cdc42 is shown to reach similar steady states in all sample simulations with the same parameter. The fluctuations are due to the dynamical mesh, a result from probabilistic Monte Carlo re-meshing, used in the simulation.

To evaluate the effect of biased cell surface material insertion, we changed the parameter used in the PDE to emulate different chemical distributions to govern the location of cell surface material insertion. We first fixed the following parameters in the chemical signal model: *k*_0_ = 20, *k*_1_ = 25, *k*_2_ = *k*_3_ = 5, *k*_4_ = 1, *k*_*ss*_ = 12, *γ* = 1, *q* = *h* = 10. In addition, *u*, which controls the bias in production of *a* is set to be *u*_*max*_ = 1.5 and *u*_*min*_ = 0.5. Instead the linear interpolation between *u*_*max*_ and *u*_*min*_ used in (5), the interpolation is made non-linear such that *u* = *u*_*min*_ + *L*_*i*_(*u*_*max*_ − *u*_*min*_) where *L*_*i*_ = (|*z*_*i*_ − *z*_*tip*_|/|*z*_*min*_ − *z*_*tip*_|)^4^ and *z, z*_*tip*_, *z*_*min*_ are the z-coordinate of the center of *i*th triangle, z-coordinate of the tip and bottom-most point, respectively. In addition, we assumed that only location with a local chemical concentration exceeding 0.8*a*_*max*_(*a*_*max*_ is the global maximum chemical concentration) is eligible for material insertion.

*β* is varied between *β* = 0.489, 0.8, 1.0 to change the size of the polarization zone such that *β* = 1.0 leading to the largest polarization zone (Fig. S4). Based on this setup, we observe different bud shapes transitioning from a relatively narrow ellipsoid to a sphere, corresponding to the increased size of the region eligible for cell surface material insertion. This suggests that the bud shape depends on where new material can be inserted, and elongation requires a highly restricted zone of material insertion or a highly polarized chemical distribution responsible for directing new materials.

**Figure S4.**
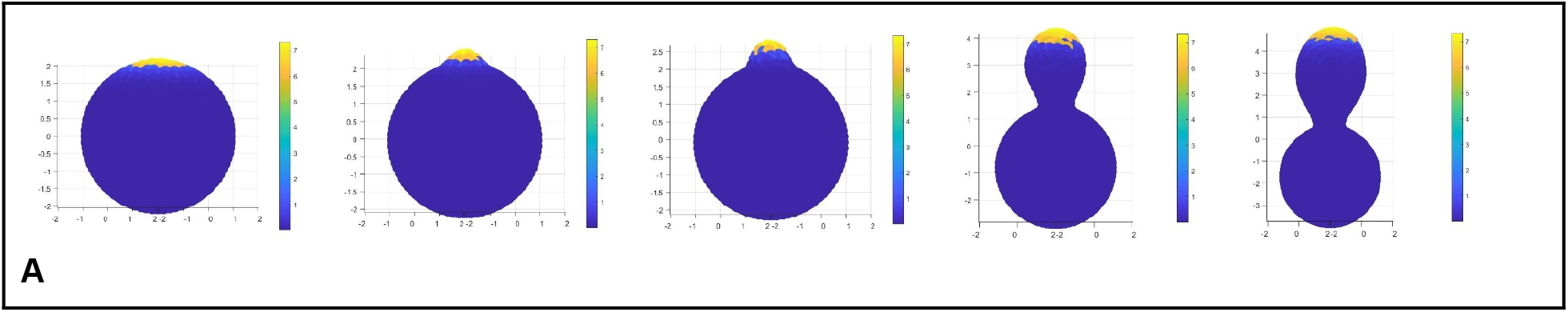
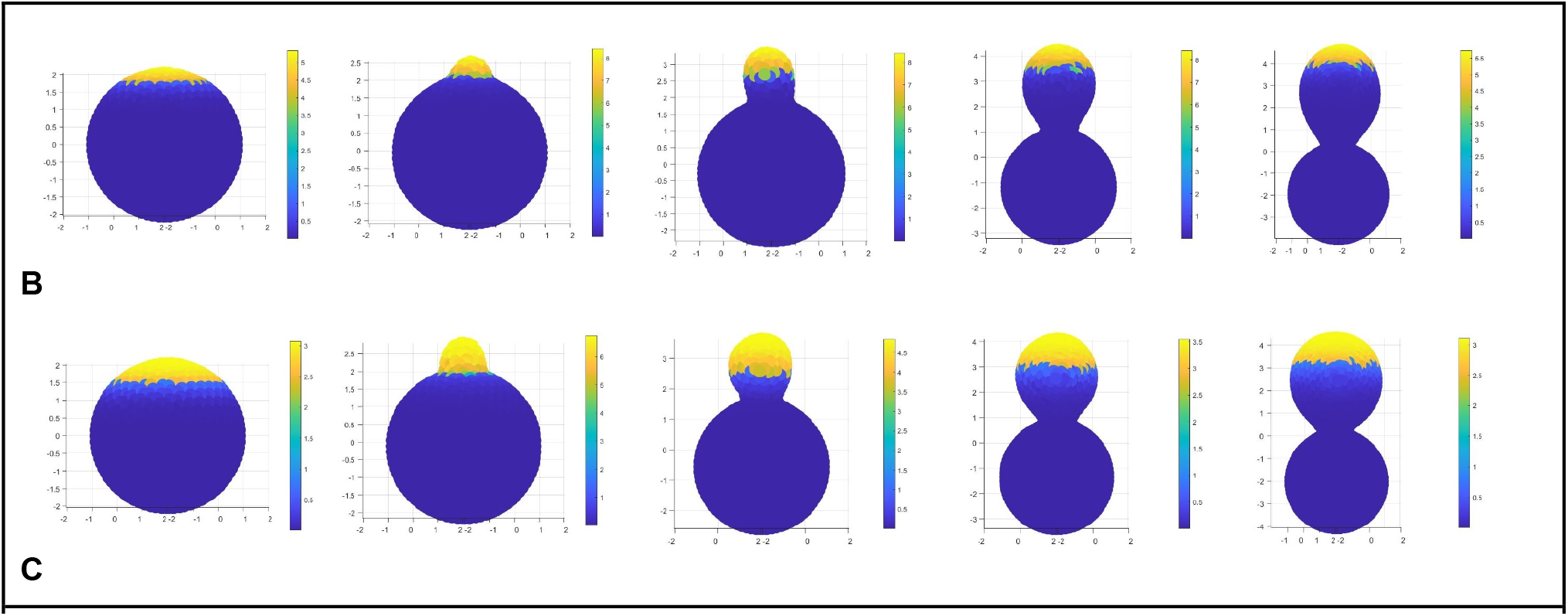
(A) *β* = 0.489, (B) *β* = 0.8, (C) *β* = 1.0. Time lapse is oriented left to right with left-most images depicting the initial polarization profiles and the right-most images depicting the final bud shape. The brightness of the color indicates the local chemical concentration effectively determining the location of material insertions. Brighter color indicates the region eligible for material insertions.

### Tubular budding cannot be generated by nonuniform mechanical properties over the bud surface within the regime of parameters required for bud initiation

It had been shown that homogeneous mechanical properties over the bud surface could only generate spherical bud shape (6). Therefore, we tested whether anisotropic mechanical properties of the bud surface were sufficient to give rise to tubular bud shape. In our previous study (6), we used a Hill function to model the anisotropic mechanical properties and tested a case that the anisotropicity occurred before the bud formation, based on the assumption that the cell polarization failed to be established uniformly within the bud region, and found that unbiological bud shapes could occur. More specifically, a “bud neck” was formed away from the septin and chitin ring position. Here again we used the Hill function to model the spatially varying mechanical properties over the bud surface with additional new assumptions. We assumed that the anisotropicity did not occur until a bud was initiated, i.e. during bud initiation, the bud surface had uniform mechanical properties. Furthermore, the midpoint between the maximally weakened mechanical properties of the bud surface and the mother cell surface was placed either at the septin and chitin ring position, or at halfway between the bud tip and the septin and chitin ring position. These new assumptions were made based on the observation that the distribution of the polarized signal which governed the budding process was uniform over the bud surface during bud initiation, but became nonuniform after the formation of a small bud (7).

**Figure S5.**
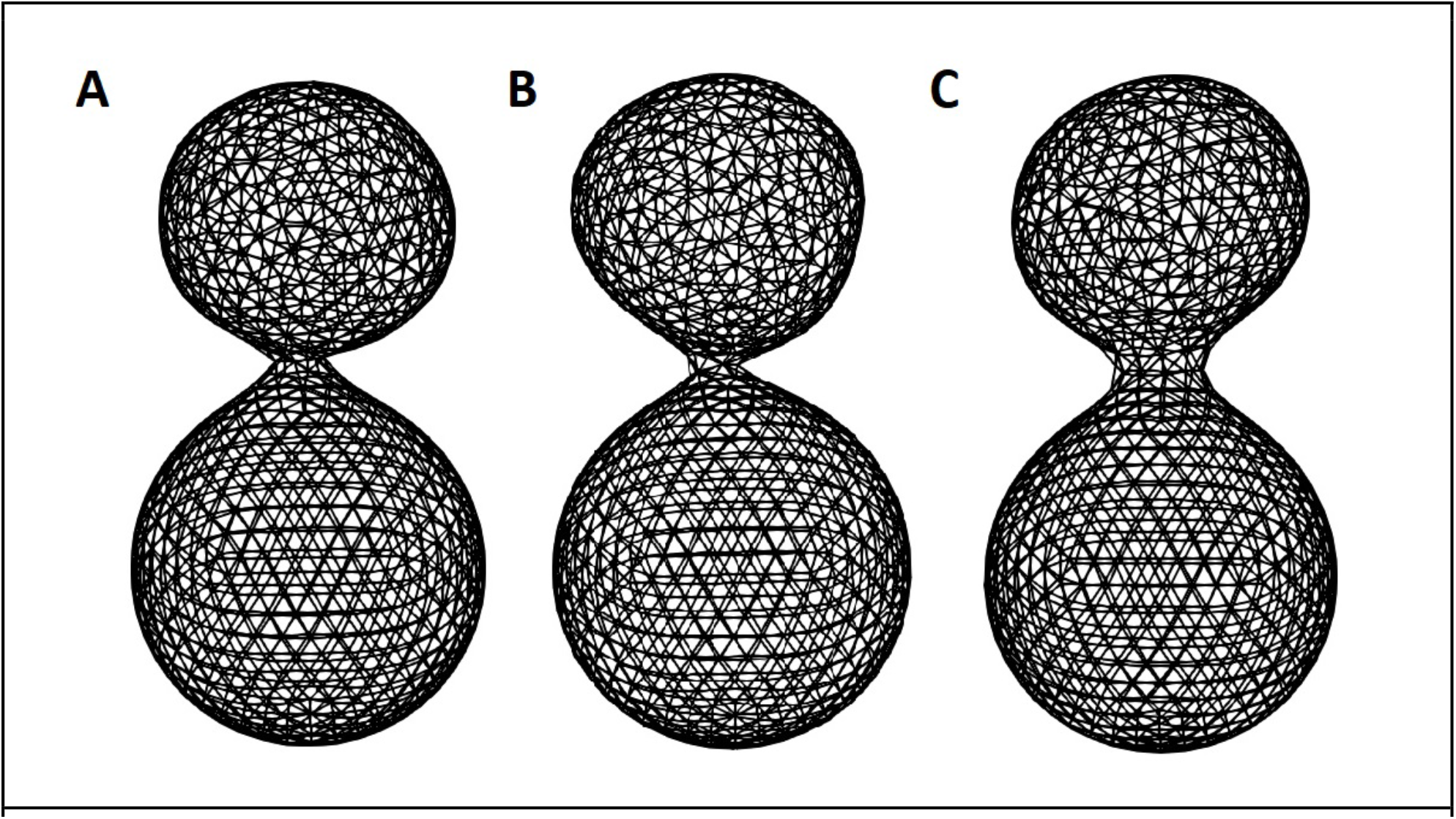
Buds with narrow neck, similar to results in previous study when the Hill function curve had a relatively sharp transition (*n* = 16). (A) Introducing the anisotropic mechanical properties from the beginning. (B) Introducing the anisotropic mechanical properties when *DR* = 4*R*_*min*_. (C) Introducing the anisotropic mechanical properties when *DR* = 6*R*_*min*_. The bud neck was more narrow if anisotropic mechanical properties were introduced earlier. Images were captured at the time point when the bud volume is about 50% of the mother cell.

In the first case such that the midpoint was at the septin and chitin ring position, the parameter sets were chosen to be (1) (*k*_*s*_, *k*_*a*_) = (1.0, 1.0) and *k*_*b*_ was varied between 0.18 and 1.0 spatially satisfying that *k*_*b*_ = 0.18 at the tip of the bud (and also uniformly at the budding site during the bud initiation) and was increased as moving toward the septin and chitin ring location, (2) *k*_*b*_ = 0.135 and *k*_*s*_, *k*_*a*_ was varied between 0.5 and 1.0 spatially satisfying that *k*_*s*_, *k*_*a*_ = 0.5 at the tip of the bud. The simulation results with high Hill coefficients still generated a “bud neck” formed away from the septin and chitin position due to the sharp change in the mechanical properties (Fig. S5), which was consistent with the results obtained in our previous study (6). Such bud formation was not sensitive to the timing of introducing the anisotropic mechanical properties which was characterized by the height of the bud obtained at that moment (termed delayed repolarization, *DR*, in our model) (Fig. S5). Shapes of buds generated in this case remained more or less spherical. In the second case such that the midpoint of the Hill function was located at the halfway between the bud tip and the septin and chitin ring position, a spherical bud similar to the experimental observation was always obtained with different Hill coefficients and different timing of introducing the anisotropic mechanical properties (Fig. S6, and Table S4, S5). Therefore, the anisotropic mechanical properties along the bud surface failed to produce tubular budding.

**Figure S6.**
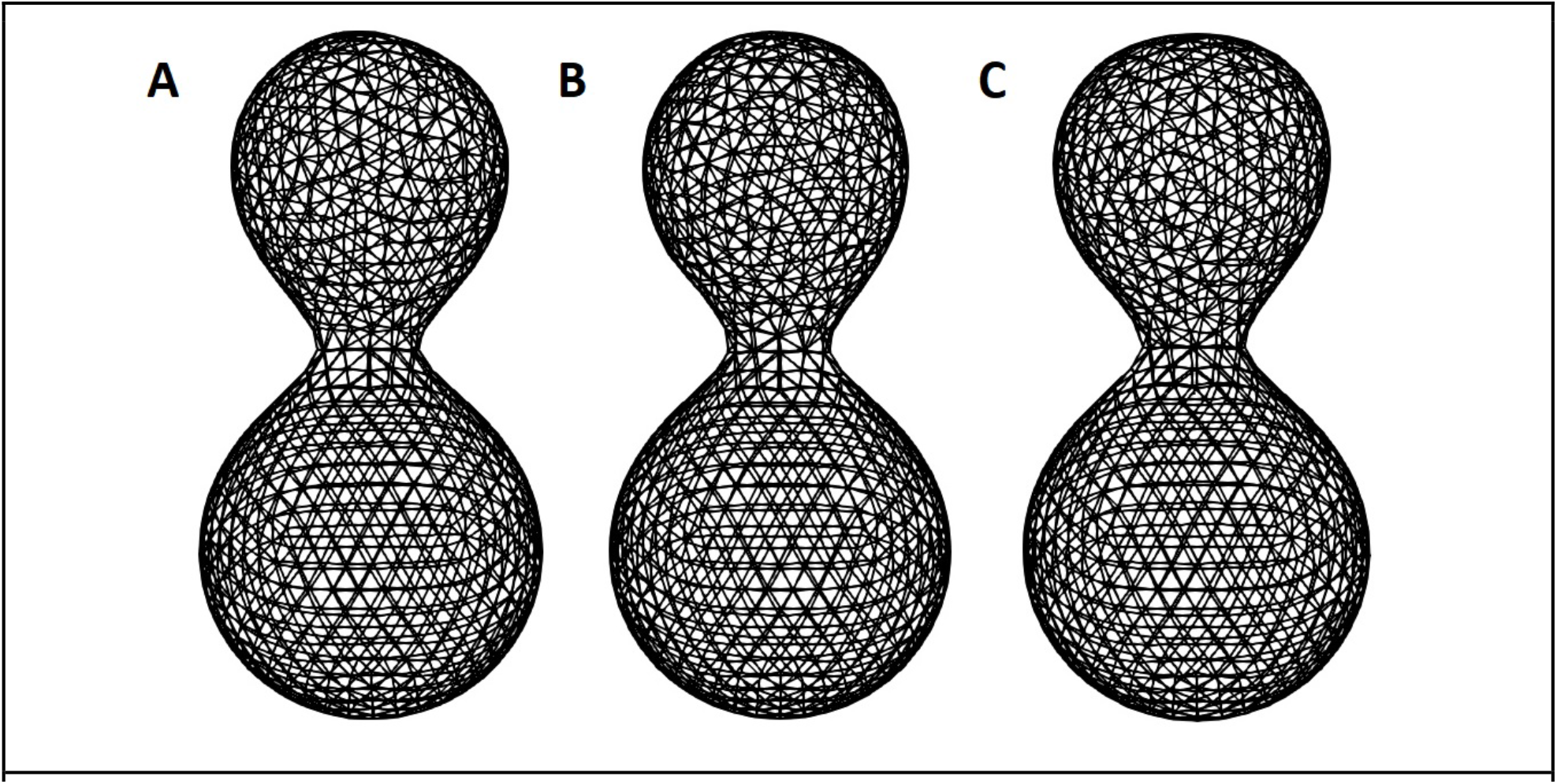
(A) *n* = 8, *K* = 0.5, *DR* = 4*R*_*min*_. (B) *n* = 8, *K* = 0.5, *DR* = 6*R*_*min*_. (C) *n* = 16, *K* = 0.5, *DR* = 4*R*_*min*_. *K* = 0.5 here indicated that the location where the Hill function yielded a value equal to the middle of its minimum and maximum was at the midpoint between the tip of the bud and the averaged z-coordinate of the septin ring. Images were captured at the time point where the total cell volume is 1.5*V*_0_. The Hill function-type scaling is applied to the bending stiffness and stretching stiffness separately.

### Perturbation of Growth Region

We assumed that new cell surface materials could only be introduced within a subregion centered at the tip of the bud instead of the entire bud. This subregion was described by the Polarization Height (PH) in our model, defined as the height of this subregion from the tip of the bud (Fig. S7A). Such an assumption was made based on the fact that chemical signals that direct the budding process polarized in different regions within the bud at different times, followed by the growth associated molecules, leading to a spatially non-homogeneous growth. Specifically, we tested *PH* = 2*R*_*min*_ corresponding to a relatively small subregion of growth and *PH* = 4*R*_*min*_ corresponding to a relatively large subregion of growth, where *R*_*min*_ is chosen to be the equilibrium length of the linear spring potential used in our model for convenience. The simulation results showed that, for *PH* = 2*R*_*min*_, a tubular budding was generated (Fig. S7A-B). The aspect ratio of the bud shape kept increasing during the growth (Fig. S7E). The relative PH to the cell height was decreasing and always at low levels, except during the bud emergence (Fig. S7F). For *PH* = 4*R*_*min*_, a spherical bud was produced with the aspect ratio maintained around 1 (Fig. S7C-D, G). The relative PH to the cell height was also decreasing over time, but it was maintained at high levels at the early stage. Although at the late stage it dropped to a similar level as the one observed for *PH* = 2*R*_*min*_ at the early stage (Fig. S7H), the spherical bud obtained at that stage was too large to change into a tubular shape. Therefore, these results together suggested that the spatially biased growth (or cell surface expansion) alone was sufficient to give rise to tubular budding once the growth region relative to the cell size was sufficiently small at the early stage of budding, even with homogeneous mechanical properties over the entire bud surface. This could be due to more restricted diffusion of the governing signaling molecules in Mode 1 due to cellular aging.

**Figure S7.**
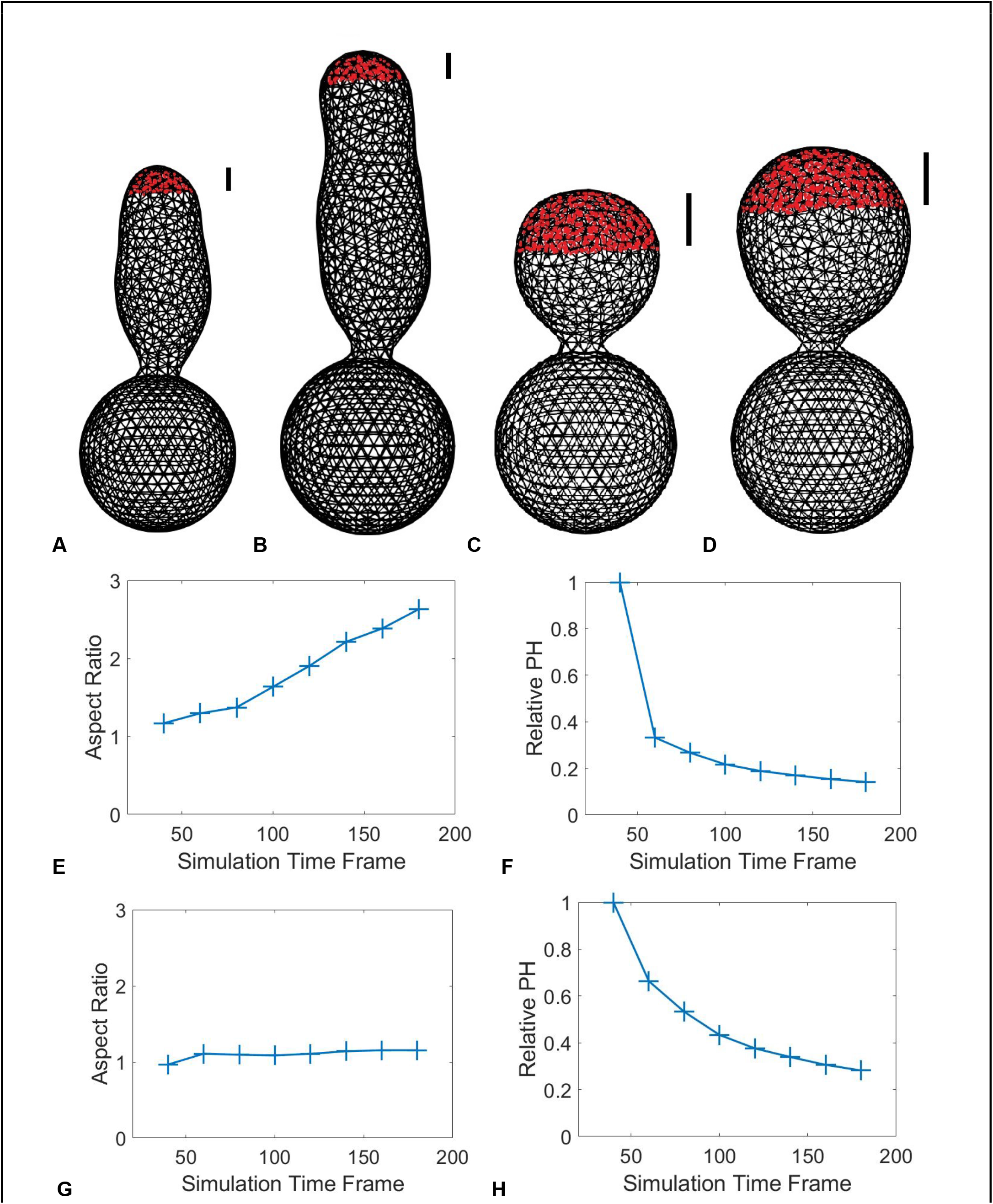
Bud shape with spatially biased growth from the beginning of budding. (A) Cell volume of 1.5*V*_0_, *PH* = 2*R*_*min*_. (B) Cell volume of 1.*76V*_0_, *PH* = 2*R*_*min*_. (C) Cell volume of 1.5*V*_0_, *PH* = 4*R*_*min*_. (D) Cell volume of 1.*97V*_0_, *PH* = 4*R*_*min*_. (E) The averaged aspect ratio of the buds produced associated with the setup in (A,B). (F) The averaged relative polarized zone height (PH) with respect to bud height associated with the setup in (A,B). (G) The averaged aspect ratio of the buds produced associated with the setup in (C,D). (H) The averaged relative PH with respect to bud height associated with the setup in (C,D). Zones in red represent the PH where new materials are allowed to be inserted for growth.

### Perturbation of Growth and Relaxation Regions

During the bud growth, enzymes involved in the cell wall modification actively cut and re-established connections among the cell wall components to allow the growth and geometric change (8). However, the exact dynamics of the cell wall modification were yet to be fully understood, and it remained unclear whether the associated enzymes were concentrated near the growth tip or acted throughout the whole bud surface. To test the effect of the spatial range of cell wall modification governed by those enzymes, we altered the region where the re-meshing technique was applied in the simulation of tubular budding with the number of re-meshing steps fixed. When the re-meshing technique was only active at the locations where new cell surface materials were introduced, the tubular bud shape could be maintained during the growth to become extremely elongated (Fig. S8A). However, since the re-meshing was restricted to the growing tip only, the bud surface exhibited some undulation outside the re-meshing region. When the re-meshing was extended further away from the bud tip, the bud became shorter and wider with less curved boundary (Fig. S8B). As we expanded the growth region, we were able to obtain a bud in a shape similar to the experimental results, which was wider at the tip and then shrunk toward the bud neck (Fig. S8C-D), though the difference due to different re-meshing regions became less significant. Overall, these simulation results suggested that the region undergoing cell wall modification also affected the bud shape. To maintain the tubular bud shape, it required less frequent cell wall modification occurring in a restricted region within the bud surface.

**Figure S8.**
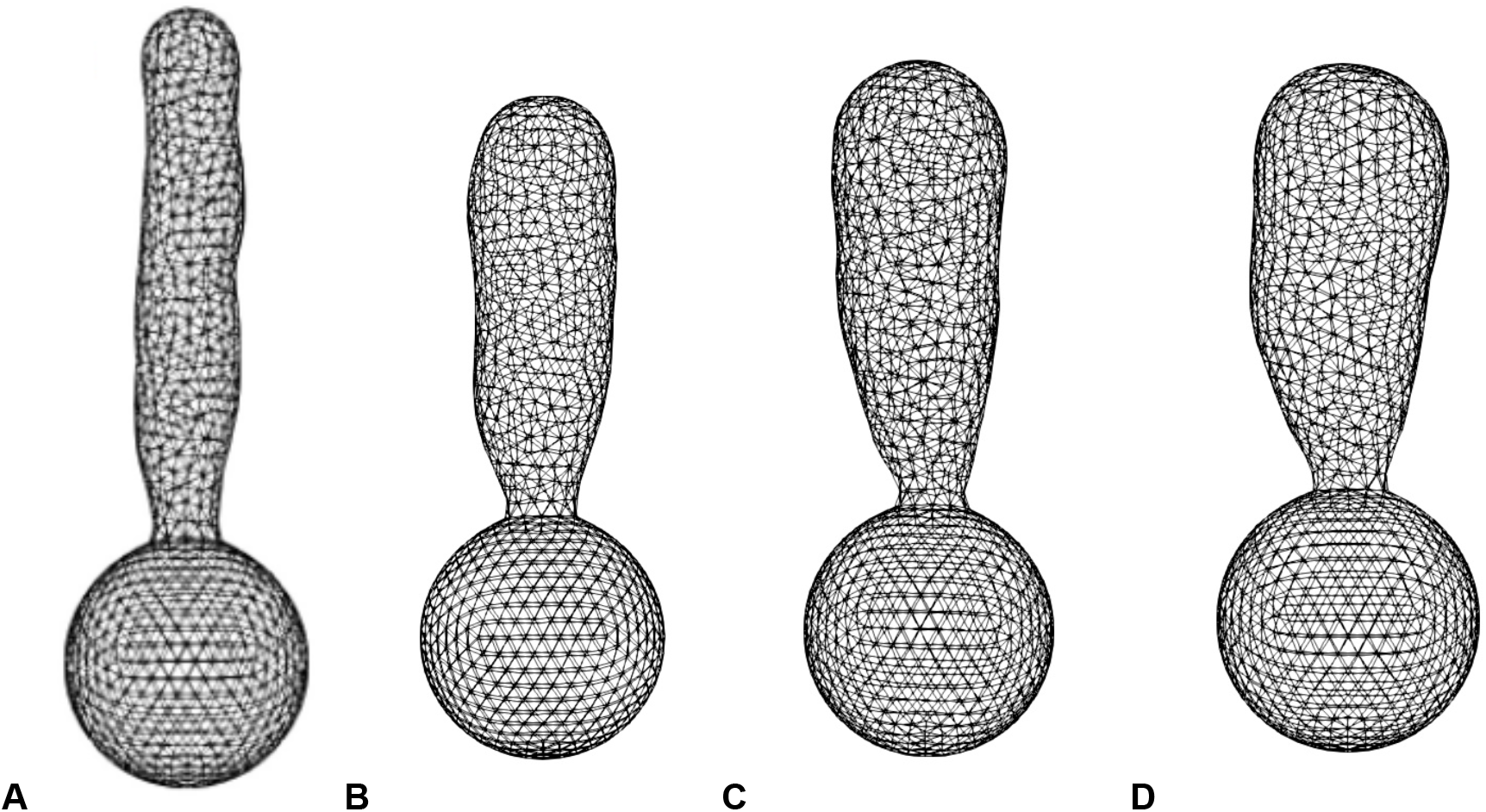
Given a fixed height for material insertion, the size of the region undergoing re-meshing can influence the final bud shape. Re-meshing and growth are applied unbiasedly throughout the whole bud until the bud heights reaches 4*R*_*min*_. Afterward, the re-meshing is restricted to regions whose height is within 2*R*_*min*_ from the bud tip (A), or restricted within 4*R*_*min*_ from the bud tip (B), whereas the material insertion is restricted within 2*R*_*min*_ from the bud tip. (C) Material insertion is restricted to 3*R*_*min*_ while the re-meshing is restricted to 2*R*_*min*_. Bud produced has an aspect ratio of 2.69. (D) Material insertion is restricted to 3*R*_*min*_ while the re-meshing is restricted to 4*R*_*min*_, and it produces a bud with lower aspect ratio, 2.20, in comparison to (C) whereas the bud volumes are similar.

### Transition from apically restricted growth to unbiased growth produces consistent bud shapes as experiments

It has been described in earlier studies that the bud shape is determined by the timing of the transition from apical growth to isotropic growth (9). To test this phenomenon in our model, we first restricted the cell growth to areas that are less than 2*L*_0_ distance (in height) away from the tip of the bud, where *L*_0_ is the equilibrium edge length used in the model. The transition from the apical growth to isotropic growth was set to occur when the height of the bud reached 10*L*_0_, 14*L*_0_, and 18*L*_0_, respectively. Once the target bud height was reached, the cell was set to grow isotropically for a fixed amount of time which accounted for 40 additional frames in the visualization output. In agreement with known facts, different transition time determined the final bud shape, i.e., 10*L*_0_, 14*L*_0_, 18*L*_0_ led to the acquisition of a spherical, a short ellipsoidal, and a long ellipsoidal bud (Fig. S9).

**Figure S9.**
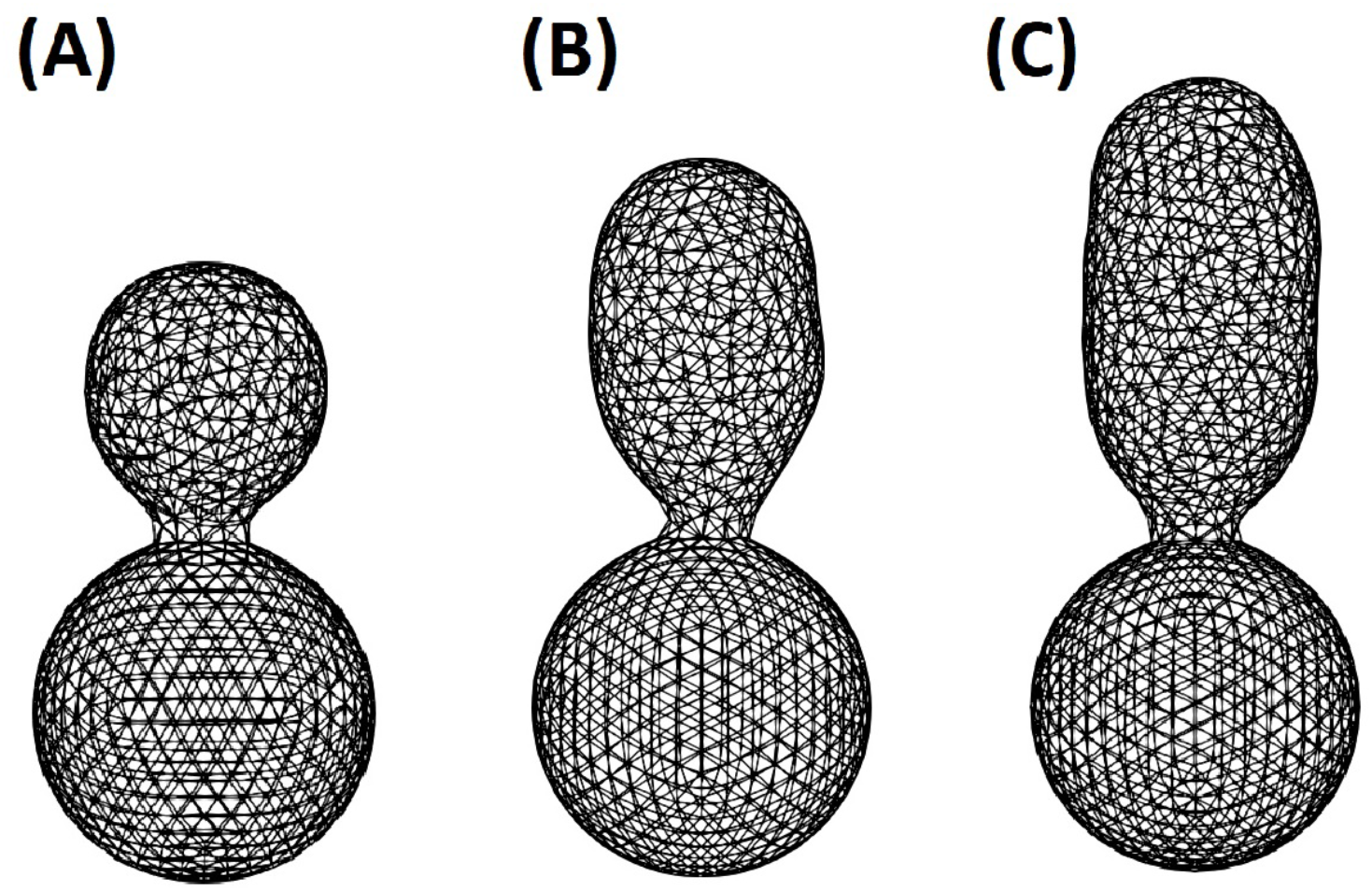
Effect of different transition time points from apical growth to isotropic growth. Transition occurred when the bud height reached (A) 10*L*_0_. (B) 14*L*_0_. (C) 18*L*_0_. Once target heights were reached, the model was set to isotropically growth for an additional 40 simulation time steps.

#### SI: Tables

**Table S1.**
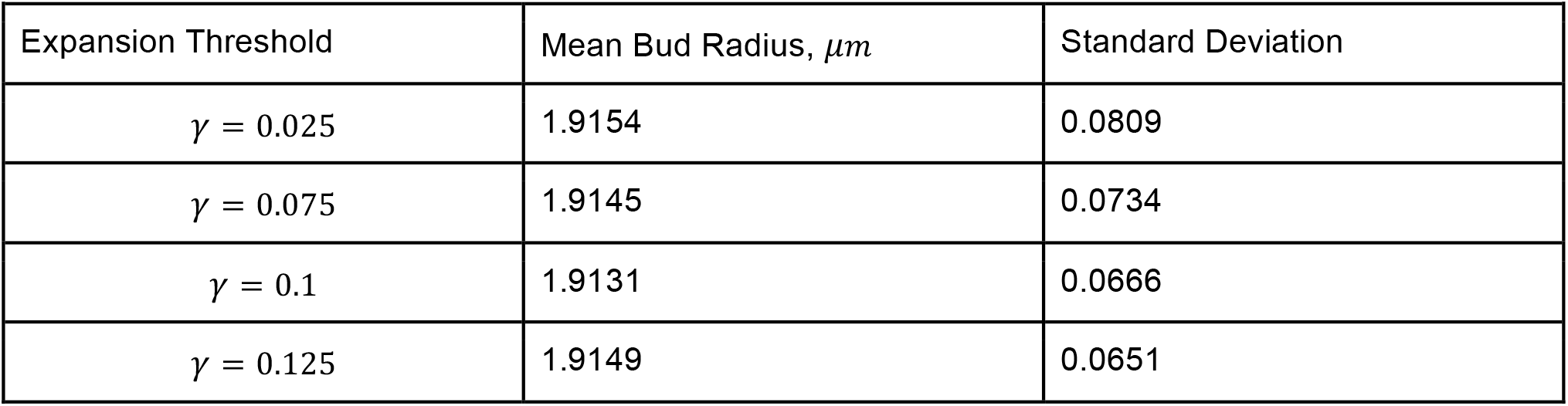
Mean bud radius and its standard deviation with respect to different expansion thresholds used.

**Table S2:**
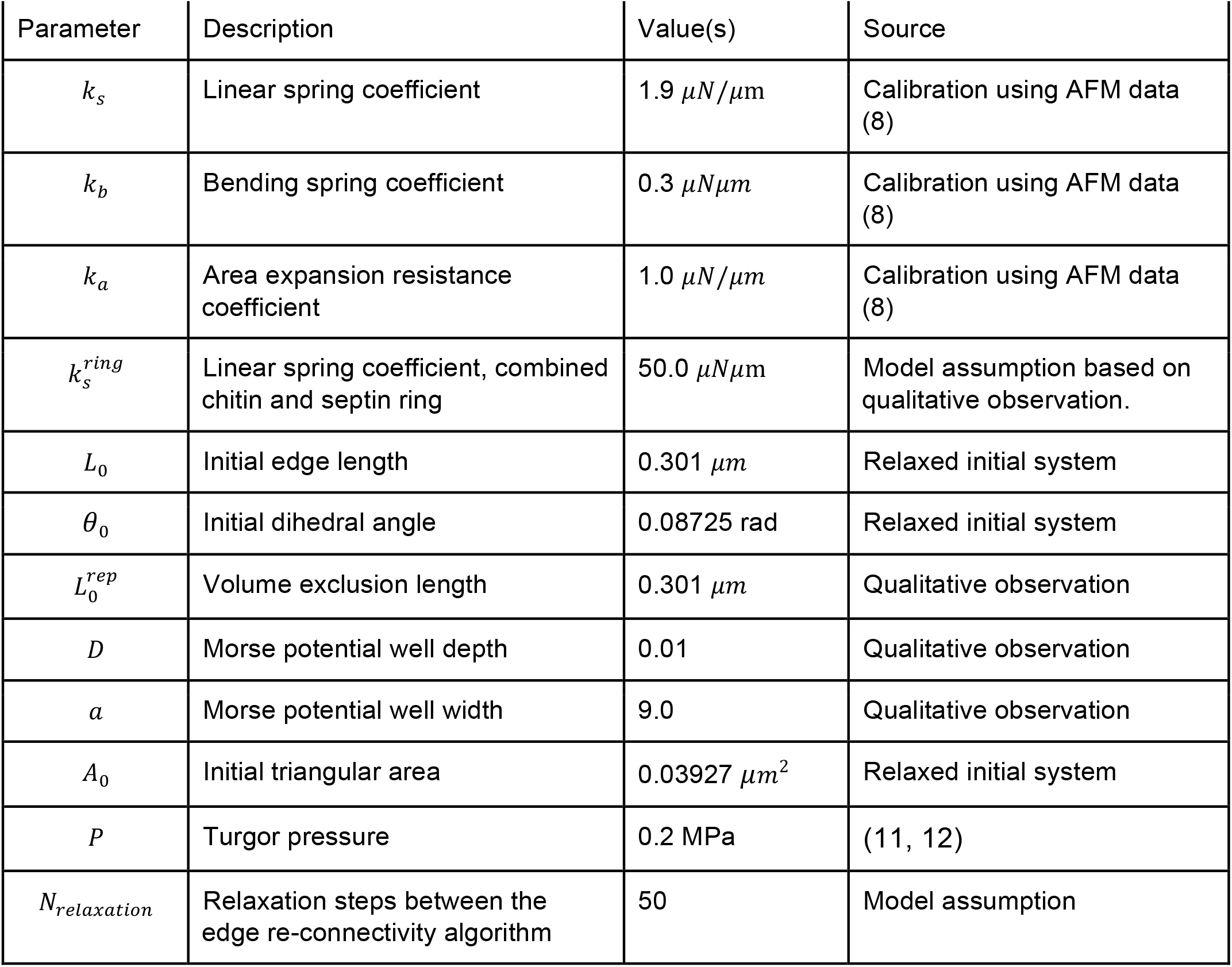
Model parameters of the yeast mother cell

**Table S3:**
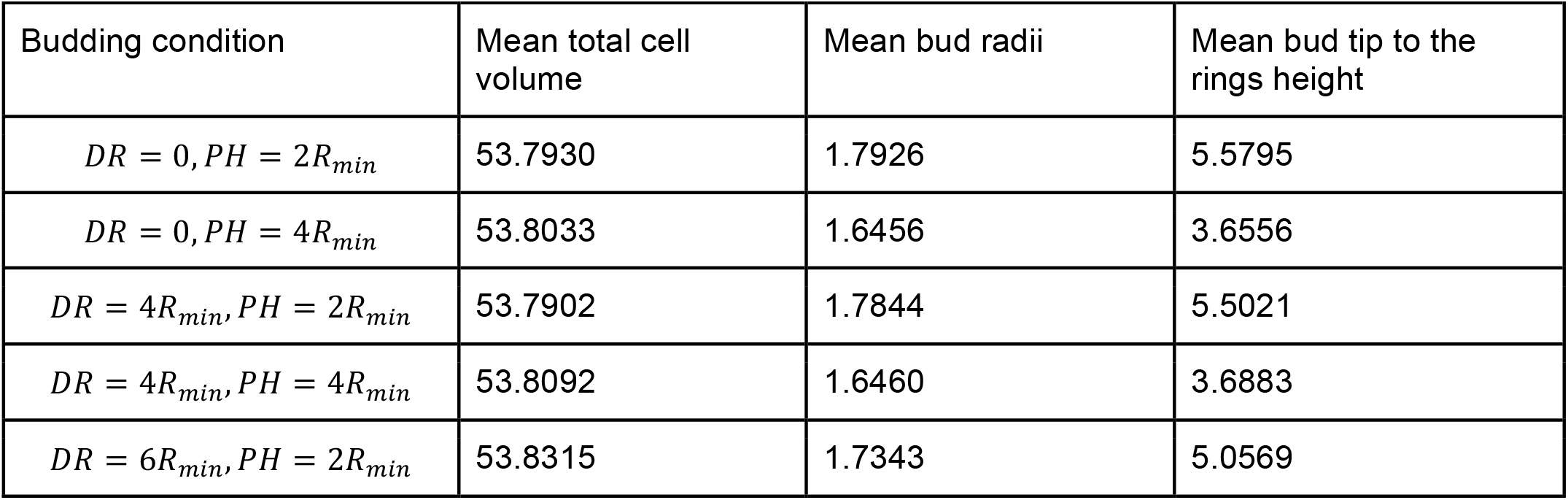
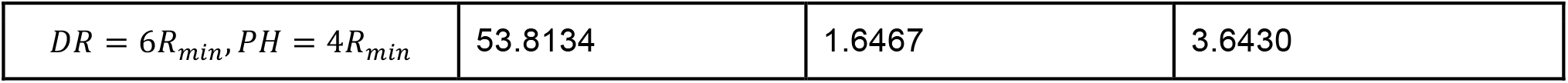
Bud radii and the height from bud tip to the septin and chitin ring positions. Delayed repolarization (DR) indicates the height of the bud to reach before the biased growth occur. Polarization height (PH) indicates the biased growth region. This region is determined by the height measured from the tip of the bud toward the mother cell. *R*_*min*_ is the equilibrium length of the linear spring potential in the model.

**Table S4:**
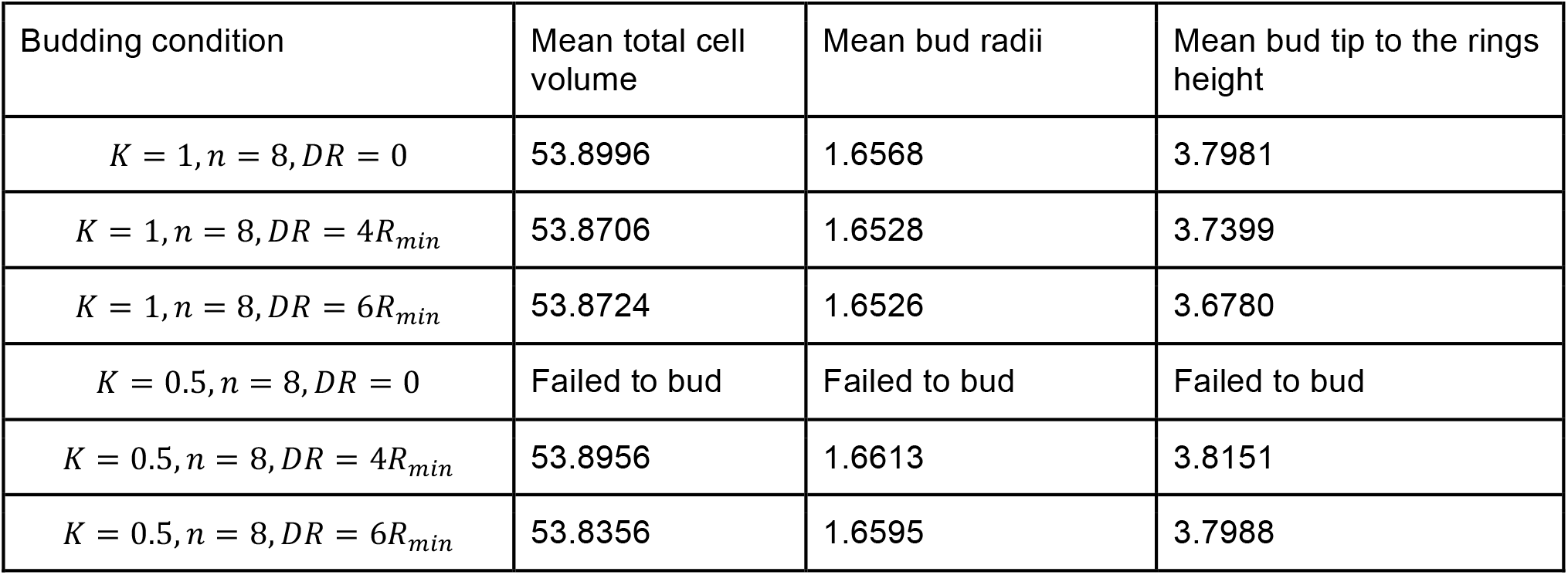
Bud radii and the height from bud tip to the septin and chitin ring positions. Delayed repolarization (DR) indicates the height of the bud to reach before anisotropic mechanical properties occur. K indicates the position of EC50 used in the Hill function such that *K* = 1 implies EC50 is chosen at the septin and chitin ring positions and *K* = 0.5 implies EC50 is chosen at the midpoint between bud tip and the ring positions. *n* is the Hill coefficient.

**Table S5:**
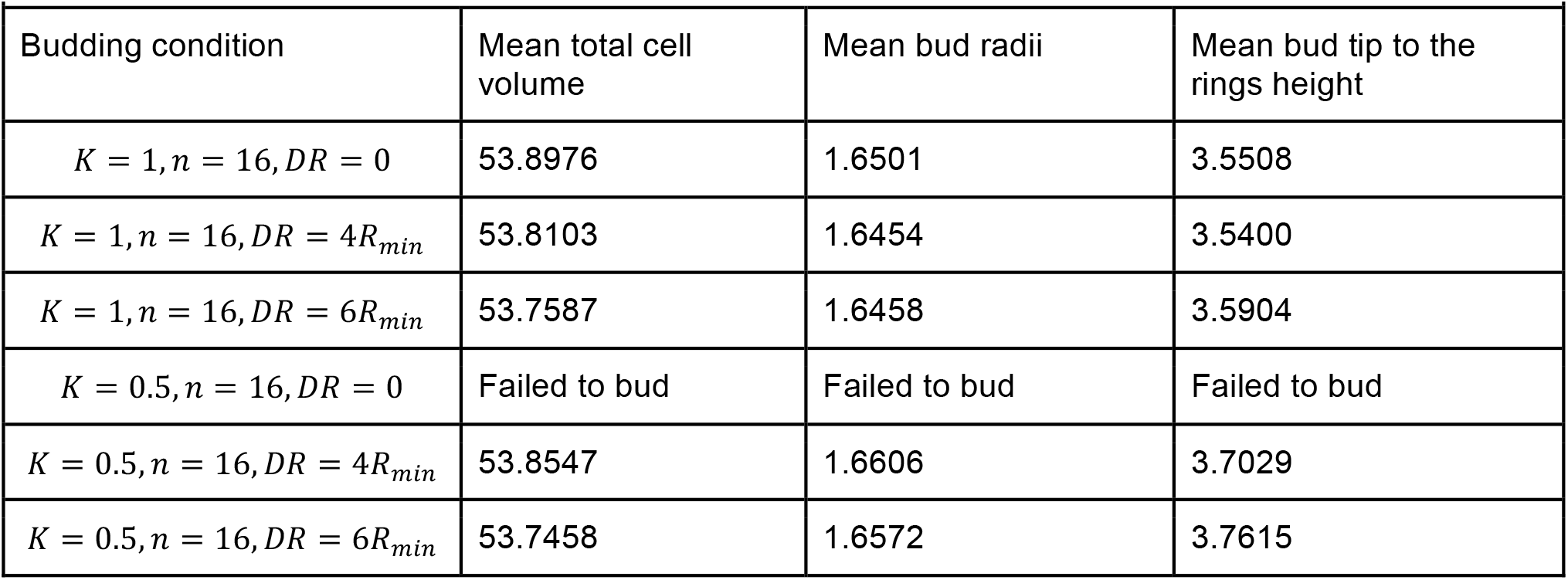
Bud radii and the height from bud tip to the septin and chitin ring positions. Delayed repolarization (DR) indicates the height of the bud to reach before anisotropic mechanical properties occur. K indicates the position of EC50 used in the Hill function such that *K* = 1 implies EC50 is chosen at the septin and chitin ring positions and *K* = 0.5 implies EC50 is chosen at the midpoint between bud tip and the ring positions. *n* is the Hill coefficient.

### Local discontinuous Galerkin (LDG) method for solving reaction-diffusion equations (1) on a surface

A LDG method developed in (9) is utilized to solve model Eqs. (1), and it is briefly described below. We first fix notations. In the present work we use a triangulated surface Γ_*h*_ composed of planar triangles *K*_*h*_ whose vertices stand on Γ to approximate cell membrane Γ. Therefore, 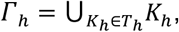, where *T*_*h*_ denotes the set of the planar triangles which form an admissible triangulation. Denote by *E* the set of edges (facets) of *T*_*h*_. For each *e* ∈ *E*, denote by *h*_*e*_ the length of the edge *e*. Let 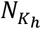 be an integer index of element *K*_*h*_, and 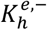 and 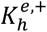 be the two elements sharing the common edge *e*. Denote by 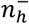 and 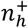 the unit outward conormal vectors defined on the edge *e* for 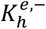 and 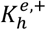, respectively. The conormal 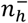 to a point *x* ∈ *e* is defined as follows according to (10) in SI References.

a. the unique unit vector that lies in the plane containing 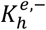;
b. 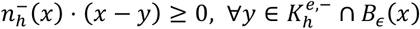, where *B*_∈_ *(x)* is a ball centered in *x*. The radius ε(> 0) of *B*_∈_ *(x)* is sufficiently small so that ∈ ≪ |*e*|, the length of edge *e*.

The conormal 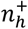 is defined similarly. With this definition, we have 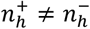 in general. Let *Pk* (*D*) denote the space of polynomials of degree not greater than *k* on any planar domain *D*. The discrete DG space *S*_*h,k*_ of scalar function associated with Γ_*h*_ is 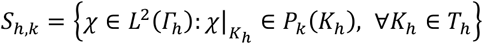, i.e., the space of piecewise polynomials which are globally in *L*^2^ (Γ_*h*_). The vector-valued DG space *Σ*_*h,k*_ associated with Γ_*h*_ is chosen to be 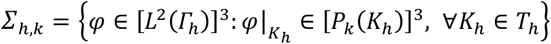. For 𝒱_*h*_ ∈ *S*_*h,k*_ and *r*_h_ ∈ *Σ*_*h,k*_, we use 𝒱 and 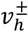 to 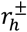 denote the trace of 𝒱_*h*_ and *r*_*h*_ on 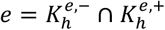 taken within the interior of 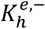 and 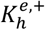, respectively. We refer readers to (13) for the definitions of the surface gradient operator ∇Γ and the laplace-Beltrami operator Δ_Γ_ on Γ. By introducing the auxiliary variable 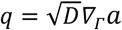, the model problem Eqs. (1) can be rewritten as a first order system of equations:

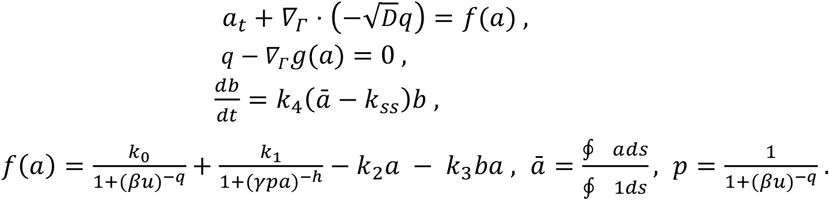

where 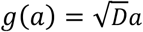. The semi-discrete LDG for solving the above equations is defined by: Find *a*_*h*_ ∈ *S*_*h,k*_ and *q*_*h*_ ∈ *Σ*_*h,k*_, such that for all test functions 𝒱_*h*_ ∈ *S*_*h,k*_ and *r*_*h*_ ∈ *Σ*_*h,k*_,

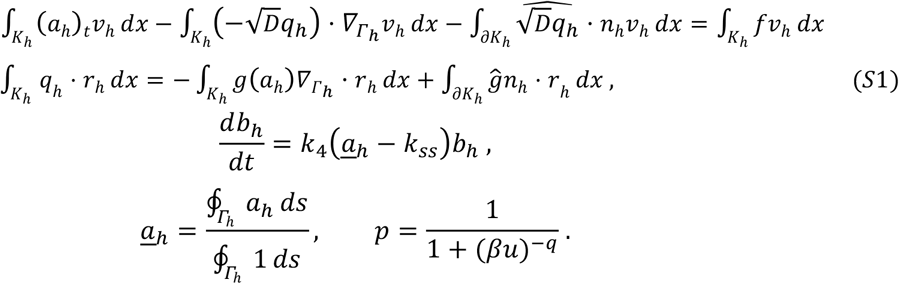

Here ĝ and 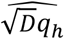 are numerical fluxes which will be described later on. To facilitate definitions of numerical fluxes, following trace operators {·} and ⟦·⟧ are introduced by following ideas in (10).

**Definition**. Denote by 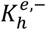 and 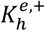 the two elements sharing the common edge *e*. For ν ∈ *L*^2^ Γ_*h*_, {ν} and ⟦ν⟧ are defined as 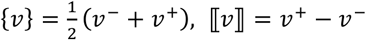 on *e*. For φ ∈ *L*^2^ Γ_*h*_ ^3^, {φ, *n*_*h*_} and ⟦φ, *n*_*h*_⟧ are defined as 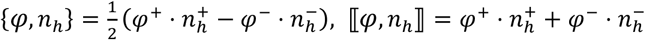 on *e*.

Denote by 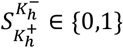 a switch function (14). 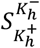 is associated with 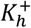 on the edge that 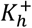 and 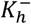 share, and is defined by:

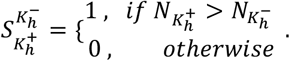

The numerical fluxes on the edge *e* for 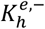 and 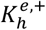 are defined respectively, as follows. The diffusive fluxes 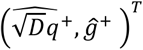 and 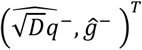:

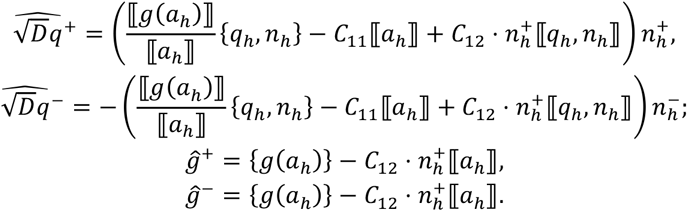

Here the penalization coefficients *C*_11_ and *C*_12_ are chosen to be

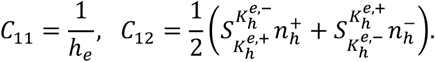

With the above choices of the numerical fluxes it yields that ⟦*ĝ*⟧ = 0 and 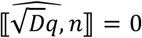. Thus these numerical fluxes are consistent and conservative. Moreover, they allow for a local resolution of *q*_2_ in terms of *a*_*h*_. The second-order accurate TVD Runge-Kutta (RK) time discretization is used to solve the semi-discrete scheme (*S*1), which can be formulated as an ordinary differential equation:

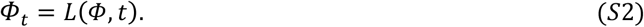

The second-order accurate TVD RK method for solving Eq. (*S*2) is given by

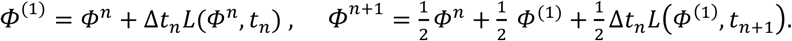

Here Δ*t*_*n*_ is the time step size. In this paper, we only considered planar triangulations of the surface which are at most second-order accurate. Therefore, we choose the second-order accurate TVD RK time-stepping method. Also, the polynomial degree *k* of the DG spaces used in this work is 1. Higher-order accurate surface approximation is needed in order to improve the overall accuracy of the scheme. We refer readers to (9) for accuracy tests of this numerical scheme.

### Movie Files

1. <u>Spherical bud</u>
2. <u>Tubular bud</u>

