## Supplementary Information for "Study of Impacts of Two Types of Cellular Aging on the Yeast Bud Morphogenesis"

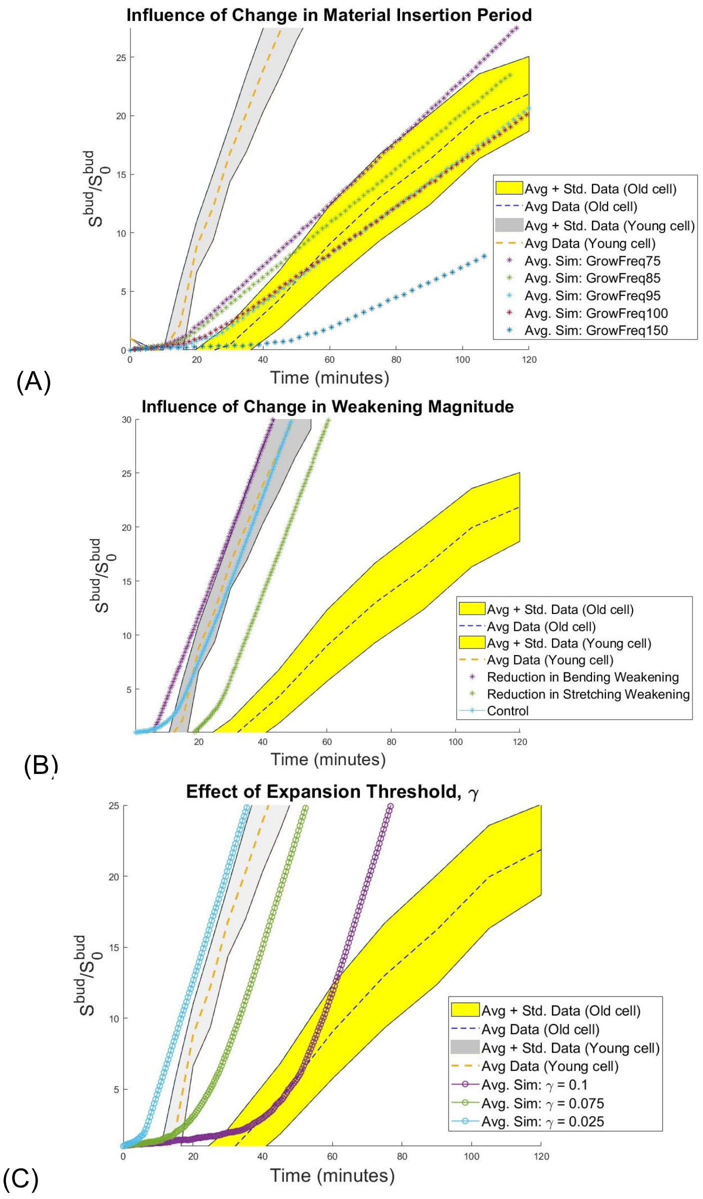


**Figure S1.** Possible changes in Mode 2 with spherical budding during cellular aging. Effect of changes in cell surface material insertion period (A) and changes in the stretching or bending stiffness (B). The y-axis represents the ratio of the bud surface area with respect to the estimated initial bud site area enclosed by the septin and chitin ring. Experimental data is collected from aged cells. (C) Effect of changes in the threshold value of surface expansion ($\gamma$). Increasing $\gamma$ led to delayed bud initiation. Bud growth trajectories showed marginal differences after bud initiation. Data presented here is extracted from young cell data.

The simulation results suggested that, for spherical budding, the growth rate was only affected by the cell surface material insertion period, whereas the bud initiation time could be affected by the mechanical properties, or the new material insertion rate, or the cell surface expansion threshold within the bud region. This indicated that the new cell surface materials were delivered less frequently to the bud region as cells aged in Mode 2. The delay of the bud initiation could be due to the less frequent new cell surface material insertion, or enhanced weakening on the mechanical properties, or a larger allocated space required for new material insertion. Therefore, appropriate parameter values were chosen in the model such that similar bud initiation time and growth rate as the experimental data could be obtained (Fig. S2).

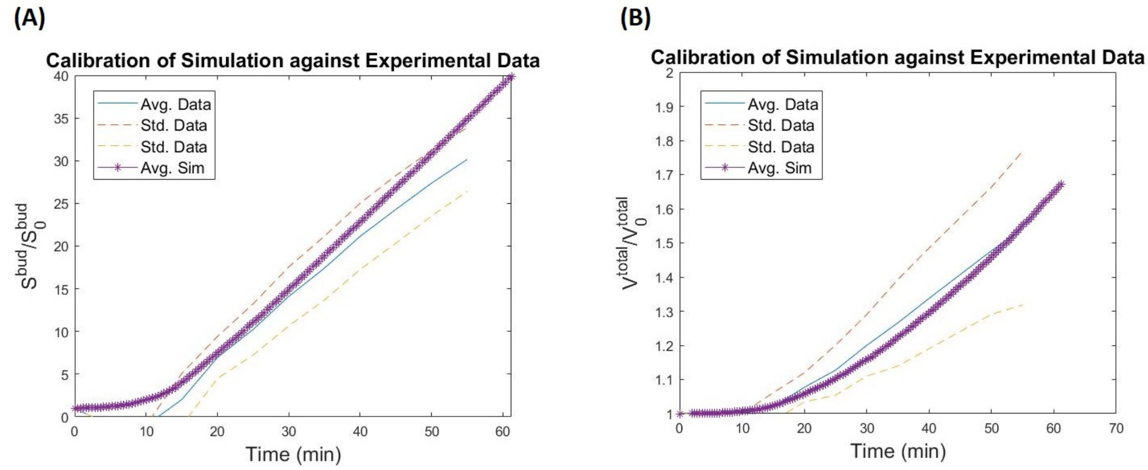


**Figure S2.** Calibration against the experimentally observed budding cycle duration for young wild type yeast cells. (A) Ratio of bud surface area with respect to initial bud surface area, and (B) Total cell volume with respect to initial total cell volume. 10 simulations were performed to ensure statistical significance. The average simulation trajectory is compared to the observed data range. Observed data range is presented as the mean trajectories with standard deviations.

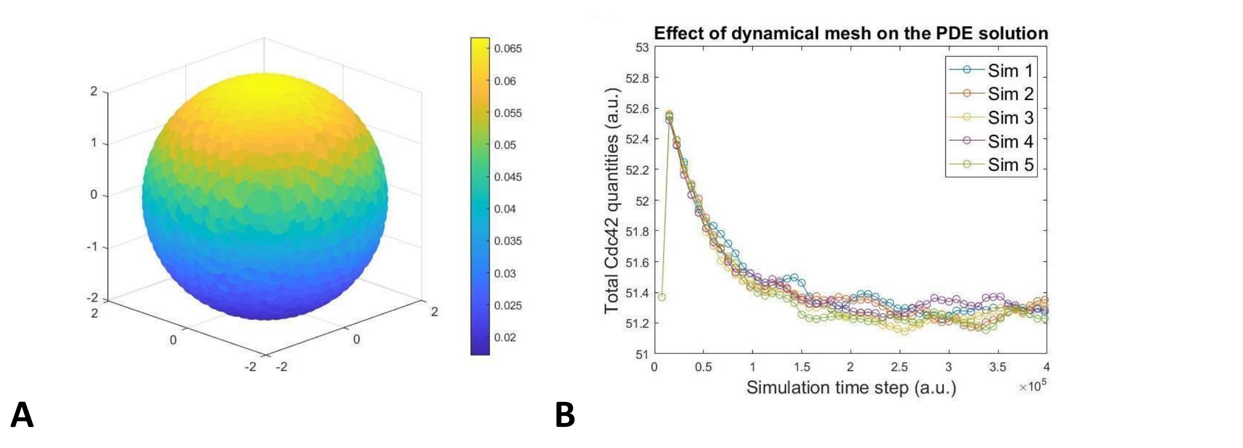


**Figure S3**. (A) A simulated polarized distribution of active Cdc42 over the triangulated sphere. The size of the polarization zone can change depending on the input parameter used for solving the differential equations. (B) The total quantity (in arbitrary units) of the Cdc42 is shown to reach similar steady states in all sample simulations with the same parameter. The fluctuations are due to the dynamical mesh, a result from probabilistic Monte Carlo re-meshing, used in the simulation.

To evaluate the effect of biased cell surface material insertion, we changed the parameter used in the PDE to emulate different chemical distributions to govern the location of cell surface material insertion. We first fixed the following parameters in the chemical signal model: $k_0 = 20, k_1 = 25, k_2 = k_3 = 5, k_4 = 1, k_{ss} = 12, \gamma = 1, q = h = 10$. In addition, $u$, which controls the bias in production of $a$ is set to be $u_{max} = 1.5$ and $u_{min} = 0.5$. Instead the linear interpolation between $u_{max}$ and $u_{min}$ used in (5), the interpolation is made non-linear such that $u = u_{min} + L_i(u_{max} - u_{min})$ where $L_i = (|z_i - z_{tip}|/|z_{min} - z_{tip}|)^4$ and $z, z_{tip}, z_{min}$ are the z-coordinate of the center of $i$th triangle, z-coordinate of the tip and bottom-most point, respectively. In addition, we assumed that only location with a local chemical concentration exceeding $0.8a_{max}$($a_{max}$ is the global maximum chemical concentration) is eligible for material insertion.

$\beta$ is varied between $\beta = 0.489, 0.8, 1.0$ to change the size of the polarization zone such that $\beta = 1.0$ leading to the largest polarization zone (Fig. S4). Based on this setup, we observe different bud shapes transitioning from a relatively narrow ellipsoid to a sphere, corresponding to the increased size of the region eligible for cell surface material insertion. This suggests that the bud shape depends on where new material can be inserted, and elongation requires a highly restricted zone of material insertion or a highly polarized chemical distribution responsible for directing new materials.

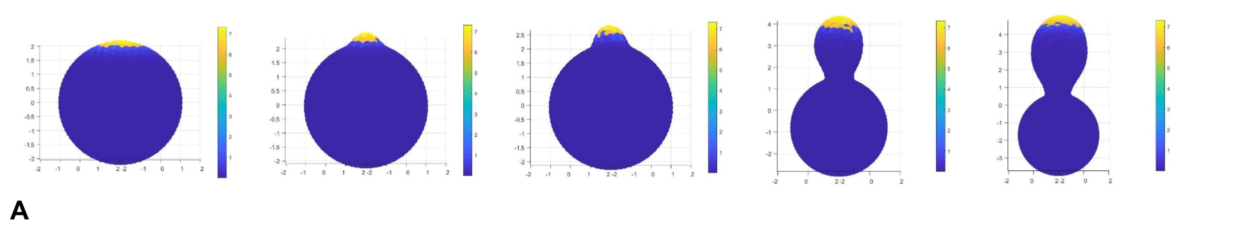

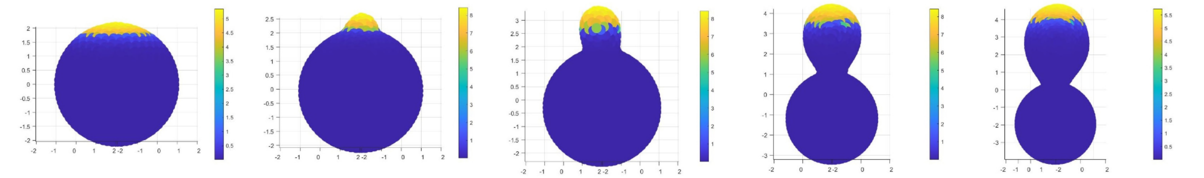

**B**

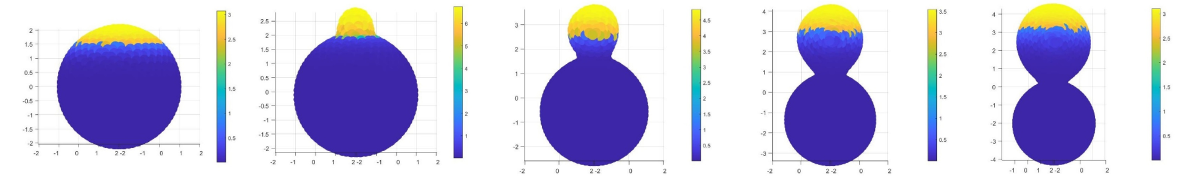

**C**

**Figure S4.** (A) $\beta = 0.489$, (B) $\beta = 0.8$, (C) $\beta = 1.0$. Time lapse is oriented left to right with left-most images depicting the initial polarization profiles and the right-most images depicting the final bud shape. The brightness of the color indicates the local chemical concentration effectively determining the location of material insertions. Brighter color indicates the region eligible for material insertions.

**Tubular budding cannot be generated by nonuniform mechanical properties over the bud surface within the regime of parameters required for bud initiation.** It had been shown that homogeneous mechanical properties over the bud surface could only generate spherical bud shape (6). Therefore, we tested whether anisotropic mechanical properties of the bud surface were sufficient to give rise to tubular bud shape. In our previous study (6), we used a Hill function to model the anisotropic mechanical properties and tested a case that the anisotropicity occurred before the bud formation, based on the assumption that the cell polarization failed to be established uniformly within the bud region, and found that unbiological bud shapes could occur. More specifically, a “bud neck” was formed away from the septin and chitin ring position. Here again we used the Hill function to model the spatially varying mechanical properties over the bud surface with additional new assumptions. We assumed that the anisotropicity did not occur until a bud was initiated, i.e. during bud initiation, the bud surface had uniform mechanical properties. Furthermore, the midpoint between the maximally weakened mechanical properties of the bud surface and the mother cell surface was placed either at the septin and chitin ring position, or at halfway between the bud tip and the septin and chitin ring position. These new assumptions were made based on the observation that the distribution of the polarized signal which governed the budding process was uniform over the bud surface during bud initiation, but became nonuniform after the formation of a small bud (7).

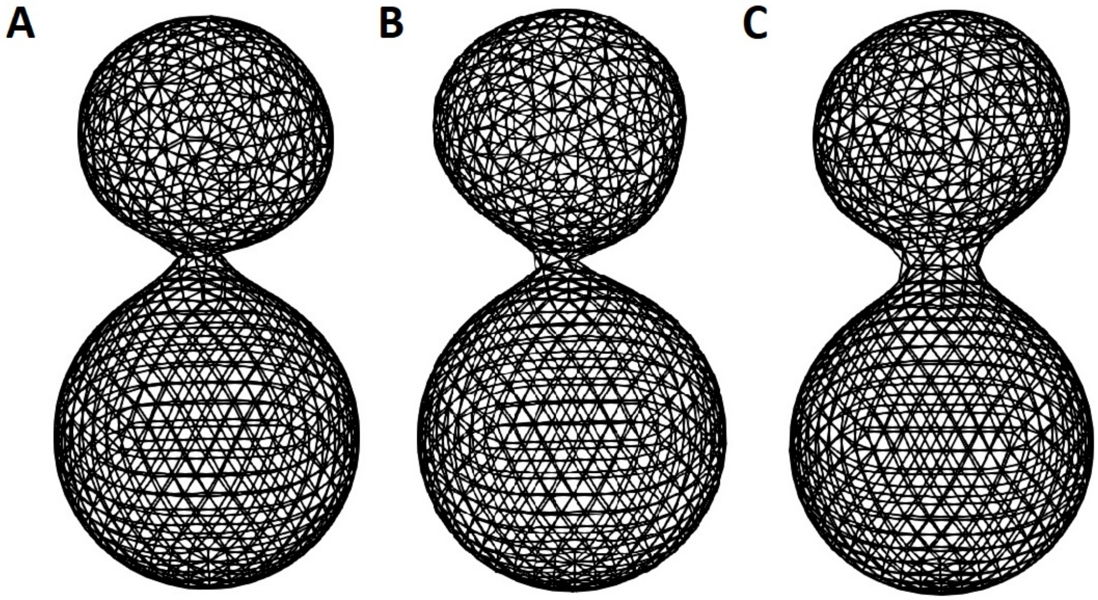


**Figure S5.** Buds with narrow neck, similar to results in previous study when the Hill function curve had a relatively sharp transition ($n = 16$). (A) Introducing the anisotropic mechanical properties from the beginning. (B) Introducing the anisotropic mechanical properties when $DR = 4R_{min}$. (C) Introducing the anisotropic mechanical properties when $DR = 6R_{min}$. The bud neck was more narrow if anisotropic mechanical properties were introduced earlier. Images were captured at the time point when the bud volume is about $50\%$ of the mother cell.

In the first case such that the midpoint was at the septin and chitin ring position, the parameter sets were chosen to be (1) $(k_s, k_a) = (1.0, 1.0)$ and $k_b$ was varied between 0.18 and 1.0 spatially satisfying that $k_b = 0.18$ at the tip of the bud (and also uniformly at the budding site during the bud initiation) and was increased as moving toward the septin and chitin ring location, (2) $k_b = 0.135$ and $k_s, k_a$ was varied between 0.5 and 1.0 spatially satisfying that $k_s, k_a = 0.5$ at the tip of the bud. The simulation results with high Hill coefficients still generated a “bud neck” formed away from the septin and chitin position due to the sharp change in the mechanical properties (Fig. S5), which was consistent with the results obtained in our previous study (6). Such bud formation was not sensitive to the timing of introducing the anisotropic mechanical properties which was characterized by the height of the bud obtained at that moment (termed delayed repolarization, $DR$, in our model) (Fig. S5). Shapes of buds generated in this case remained more or less spherical. In the second case such that the midpoint of the Hill function was located at the halfway between the bud tip and the septin and chitin ring position, a spherical bud similar to the experimental observation was always obtained with different Hill coefficients and different timing of introducing the anisotropic mechanical properties (Fig. S6, and Table S4, S5). Therefore, the anisotropic mechanical properties along the bud surface failed to produce tubular budding.

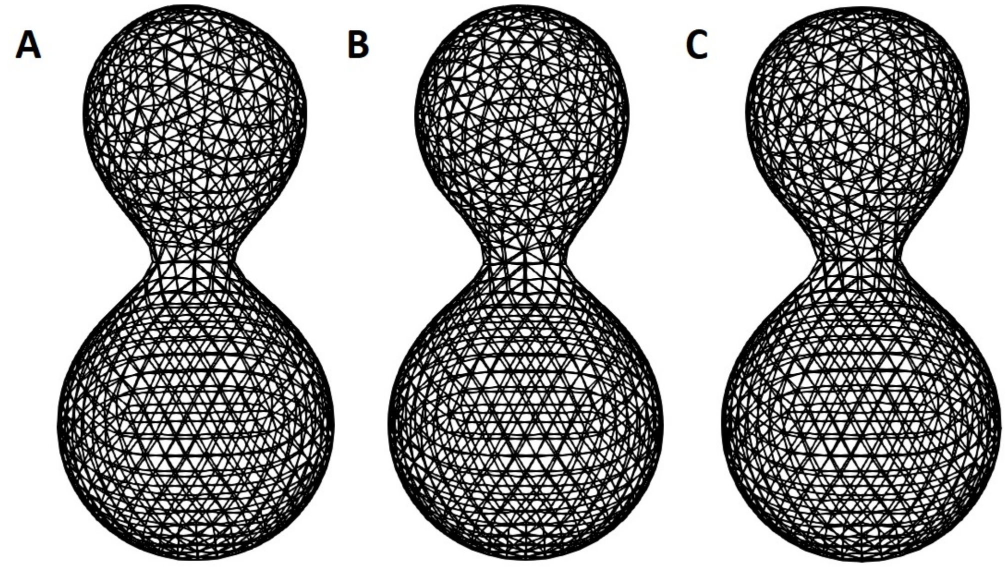


**Figure S6.** (A) $n = 8, K = 0.5, DR = 4R_{min}$. (B) $n = 8, K = 0.5, DR = 6R_{min}$. (C) $n = 16, K = 0.5, DR = 4R_{min}$. $K = 0.5$ here indicated that the location where the Hill function yielded a value equal to the middle of its minimum and maximum was at the midpoint between the tip of the bud and the averaged z-coordinate of the septin ring. Images were captured at the time point where the total cell volume is $1.5V_0$. The Hill function-type scaling is applied to the bending stiffness and stretching stiffness separately.

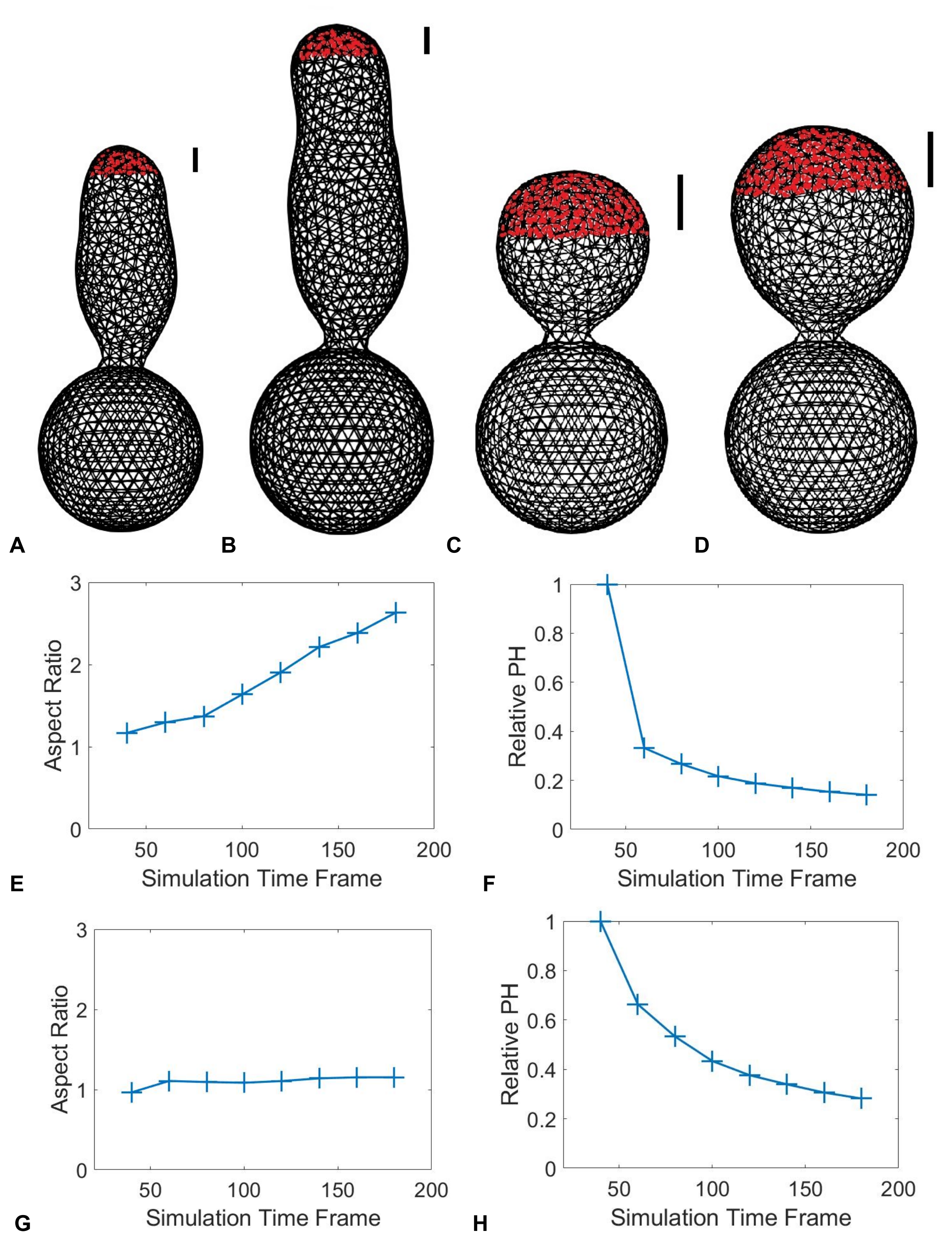


**Figure S7.** Bud shape with spatially biased growth from the beginning of budding. (A) Cell volume of $1.5V_0$, $PH = 2R_{min}$. (B) Cell volume of $1.76V_0$, $PH = 2R_{min}$. (C) Cell volume of $1.5V_0$, $PH = 4R_{min}$. (D) Cell volume of $1.97V_0$, $PH = 4R_{min}$. (E) The averaged aspect ratio of the buds produced associated with the setup in (A,B). (F) The averaged relative polarized zone height (PH) with respect to bud height associated with the setup in (A,B). (G) The averaged aspect ratio of the buds produced associated with the setup in (C,D). (H) The averaged relative PH with respect to bud height associated with the setup in (C,D). Zones in red represent the PH where new materials are allowed to be inserted for growth.

**Perturbation of Growth and Relaxation Regions.** During the bud growth, enzymes involved in the cell wall modification actively cut and re-established connections among the cell wall components to allow the growth and geometric change (8). However, the exact dynamics of the cell wall modification were yet to be fully understood, and it remained unclear whether the associated enzymes were concentrated near the growth tip or acted throughout the whole bud surface. To test the effect of the spatial range of cell wall modification governed by those enzymes, we altered the region where the re-meshing technique was applied in the simulation of tubular budding with the number of re-meshing steps fixed. When the re-meshing technique was only active at the locations where new cell surface materials were introduced, the tubular bud shape could be maintained during the growth to become extremely elongated (Fig. S8A). However, since the re-meshing was restricted to the growing tip only, the bud surface exhibited some undulation outside the re-meshing region. When the re-meshing was extended further away from the bud tip, the bud became shorter and wider with less curved boundary (Fig. S8B). As we expanded the growth region, we were able to obtain a bud in a shape similar to the experimental results, which was wider at the tip and then shrunk toward the bud neck (Fig. S8C-D), though the difference due to different re-meshing regions became less significant. Overall, these simulation results suggested that the region undergoing cell wall modification also affected the bud shape. To maintain the tubular bud shape, it required less frequent cell wall modification occurring in a restricted region within the bud surface.

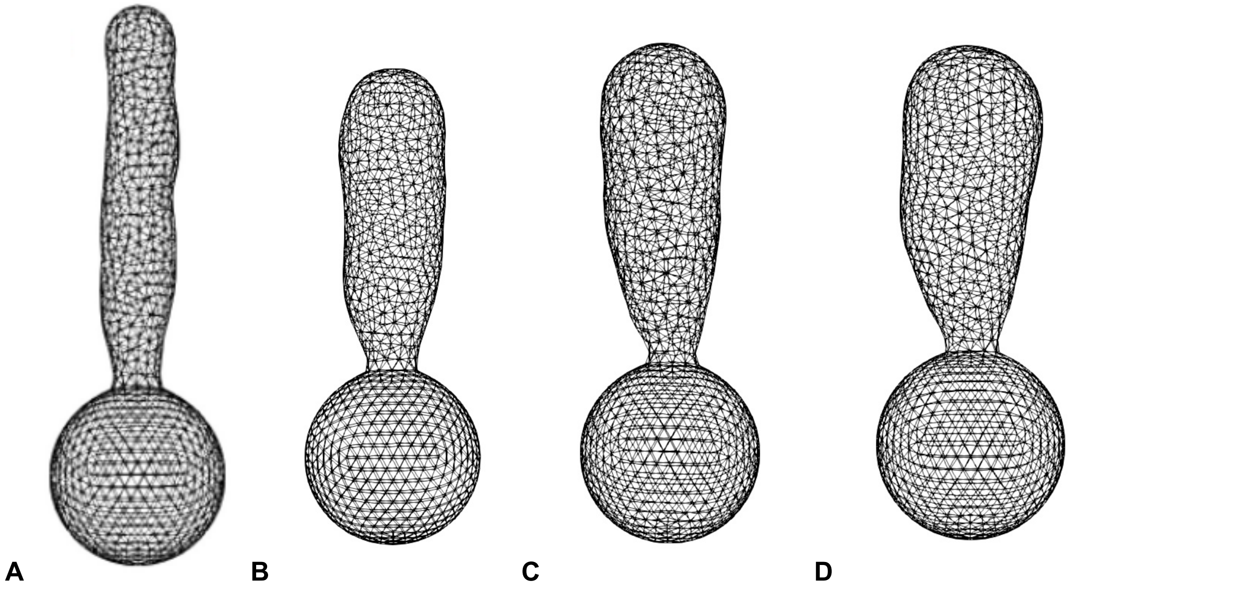


**Figure S8**. Given a fixed height for material insertion, the size of the region undergoing re-meshing can influence the final bud shape. Re-meshing and growth are applied unbiasedly throughout the whole bud until the bud heights reaches $4R_{min}$. Afterward, the re-meshing is restricted to regions whose height is within $2R_{min}$ from the bud tip (A), or restricted within $4R_{min}$ from the bud tip (B), whereas the material insertion is restricted within $2R_{min}$ from the bud tip. (C) Material insertion is restricted to $3R_{min}$ while the re-meshing is restricted to $2R_{min}$. Bud produced has an aspect ratio of 2.69. (D) Material insertion is restricted to $3R_{min}$ while the re-meshing is restricted to $4R_{min}$, and it produces a bud with lower aspect ratio, 2.20, in comparison to (C) whereas the bud volumes are similar.

**Transition from apically restricted growth to unbiased growth produces consistent bud shapes as experiments.** It has been described in earlier studies that the bud shape is determined by the timing of the transition from apical growth to isotropic growth (9). To test this phenomenon in our model, we first restricted the cell growth to areas that are less than $2L_0$ distance (in height) away from the tip of the bud, where $L_0$ is the equilibrium edge length used in the model. The transition from the apical growth to isotropic growth was set to occur when the height of the bud reached $10L_0, 14L_0$, and $18L_0$, respectively. Once the target bud height was reached, the cell was set to grow isotropically for a fixed amount of time which accounted for 40 additional frames in the visualization output. In agreement with known facts, different transition time determined the final bud shape, i.e., $10L_0, 14L_0, 18L_0$ led to the acquisition of a spherical, a short ellipsoidal, and a long ellipsoidal bud (Fig. S9).

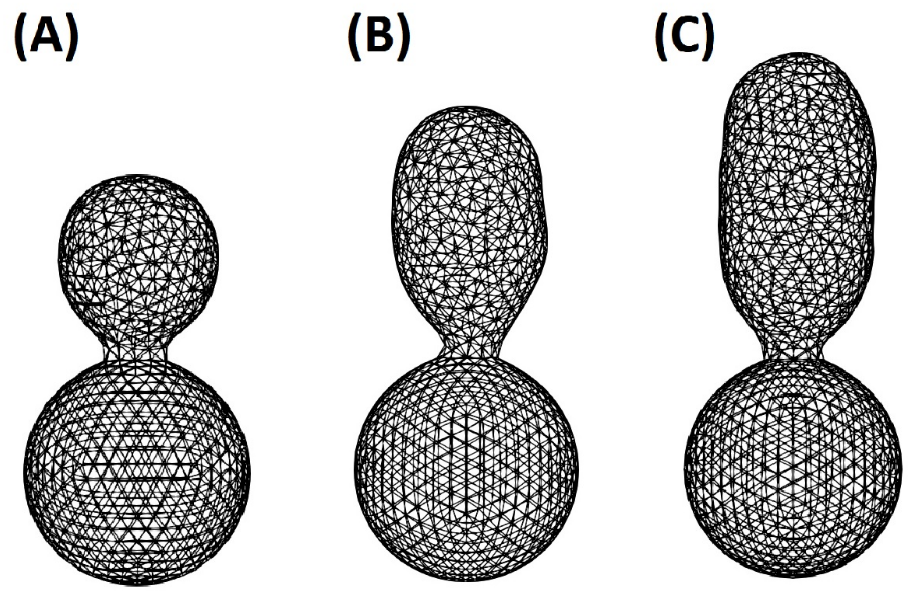


**Figure S9**. Effect of different transition time points from apical growth to isotropic growth. Transition occurred when the bud height reached (A) $10L_0$. (B) $14L_0$. (C) $18L_0$. Once target heights were reached, the model was set to isotropically growth for an additional 40 simulation time steps.

**SI: Tables**

**Table S1**. Mean bud radius and its standard deviation with respect to different expansion thresholds used.

| Expansion Threshold | Mean Bud Radius, $\mu m$ | Standard Deviation |
|---|---|---|
| $\gamma = 0.025$ | 1.9154 | 0.0809 |
| $\gamma = 0.075$ | 1.9145 | 0.0734 |
| $\gamma = 0.1$ | 1.9131 | 0.0666 |
| $\gamma = 0.125$ | 1.9149 | 0.0651 |

| **Table S2:** Model parameters of the yeast mother cell | | | |
|---|---|---|---|
| Parameter | Description | Value(s) | Source |
| $k_s$ | Linear spring coefficient | 1.9 $\mu N/\mu$m | Calibration using AFM data (8) |
| $k_b$ | Bending spring coefficient | 0.3 $\mu N \mu m$ | Calibration using AFM data (8) |
| $k_a$ | Area expansion resistance coefficient | 1.0 $\mu N/\mu m$ | Calibration using AFM data (8) |
| $k_s^{ring}$ | Linear spring coefficient, combined chitin and septin ring | 50.0 $\mu N \mu$m | Model assumption based on qualitative observation. |
| $L_0$ | Initial edge length | 0.301 $\mu m$ | Relaxed initial system |
| $\theta_0$ | Initial dihedral angle | 0.08725 rad | Relaxed initial system |
| $L_0^{rep}$ | Volume exclusion length | 0.301 $\mu m$ | Qualitative observation |
| $D$ | Morse potential well depth | 0.01 | Qualitative observation |
| $a$ | Morse potential well width | 9.0 | Qualitative observation |
| $A_0$ | Initial triangular area | 0.03927 $\mu m^2$ | Relaxed initial system |
| $P$ | Turgor pressure | 0.2 MPa | (11, 12) |
| $N_{relaxation}$ | Relaxation steps between the edge re-connectivity algorithm | 50 | Model assumption |

| **Table S3**: Bud radii and the height from bud tip to the septin and chitin ring positions. Delayed repolarization (DR) indicates the height of the bud to reach before the biased growth occur. Polarization height (PH) indicates the biased growth region. This region is determined by the height measured from the tip of the bud toward the mother cell. $R_{min}$ is the equilibrium length of the linear spring potential in the model. | | | |
|---|---|---|---|
| Budding condition | Mean total cell volume | Mean bud radii | Mean bud tip to the rings height |
| $DR = 0, PH = 2R_{min}$ | 53.7930 | 1.7926 | 5.5795 |
| $DR = 0, PH = 4R_{min}$ | 53.8033 | 1.6456 | 3.6556 |
| $DR = 4R_{min}, PH = 2R_{min}$ | 53.7902 | 1.7844 | 5.5021 |
| $DR = 4R_{min}, PH = 4R_{min}$ | 53.8092 | 1.6460 | 3.6883 |
| $DR = 6R_{min}, PH = 2R_{min}$ | 53.8315 | 1.7343 | 5.0569 |

| | | | |
|---|---|---|---|
| $DR = 6R_{min}, PH = 4R_{min}$ | 53.8134 | 1.6467 | 3.6430 |

**Table S4**: Bud radii and the height from bud tip to the septin and chitin ring positions. Delayed repolarization (DR) indicates the height of the bud to reach before anisotropic mechanical properties occur. K indicates the position of EC50 used in the Hill function such that $K = 1$ implies EC50 is chosen at the septin and chitin ring positions and $K = 0.5$ implies EC50 is chosen at the midpoint between bud tip and the ring positions. $n$ is the Hill coefficient.

| Budding condition | Mean total cell volume | Mean bud radii | Mean bud tip to the rings height |
|---|---|---|---|
| $K = 1, n = 8, DR = 0$ | 53.8996 | 1.6568 | 3.7981 |
| $K = 1, n = 8, DR = 4R_{min}$ | 53.8706 | 1.6528 | 3.7399 |
| $K = 1, n = 8, DR = 6R_{min}$ | 53.8724 | 1.6526 | 3.6780 |
| $K = 0.5, n = 8, DR = 0$ | Failed to bud | Failed to bud | Failed to bud |
| $K = 0.5, n = 8, DR = 4R_{min}$ | 53.8956 | 1.6613 | 3.8151 |
| $K = 0.5, n = 8, DR = 6R_{min}$ | 53.8356 | 1.6595 | 3.7988 |

**Table S5**: Bud radii and the height from bud tip to the septin and chitin ring positions. Delayed repolarization (DR) indicates the height of the bud to reach before anisotropic mechanical properties occur. K indicates the position of EC50 used in the Hill function such that $K = 1$ implies EC50 is chosen at the septin and chitin ring positions and $K = 0.5$ implies EC50 is chosen at the midpoint between bud tip and the ring positions. $n$ is the Hill coefficient.

| Budding condition | Mean total cell volume | Mean bud radii | Mean bud tip to the rings height |
|---|---|---|---|
| $K = 1, n = 16, DR = 0$ | 53.8976 | 1.6501 | 3.5508 |
| $K = 1, n = 16, DR = 4R_{min}$ | 53.8103 | 1.6454 | 3.5400 |
| $K = 1, n = 16, DR = 6R_{min}$ | 53.7587 | 1.6458 | 3.5904 |
| $K = 0.5, n = 16, DR = 0$ | Failed to bud | Failed to bud | Failed to bud |
| $K = 0.5, n = 16, DR = 4R_{min}$ | 53.8547 | 1.6606 | 3.7029 |
| $K = 0.5, n = 16, DR = 6R_{min}$ | 53.7458 | 1.6572 | 3.7615 |

**Local discontinuous Galerkin (LDG) method for solving reaction-diffusion equations (1) on a surface.** A LDG method developed in (9) is utilized to solve model Eqs. (1), and it is briefly described below. We first fix notations. In the present work we use a triangulated surface $\Gamma_h$ composed of planar triangles $K_h$ whose vertices stand on $\Gamma$ to approximate cell membrane $\Gamma$. Therefore, $\Gamma_h = \bigcup_{K_h \in T_h} K_h$, where $T_h$ denotes the set of the planar triangles which form an admissible triangulation. Denote by $E$ the set of edges (facets) of $T_h$. For each $e \in E$, denote by $h_e$ the length of the edge $e$. Let $N_{K_h}$ be an integer index of element $K_h$, and $K_h^{e,-}$ and $K_h^{e,+}$ be the two elements sharing the common edge $e$. Denote by $n_h^-$

and $n_h^+$ the unit outward conormal vectors defined on the edge $e$ for $K_h^{e,-}$ and $K_h^{e,+}$, respectively. The conormal $n_h^-$ to a point $x \in e$ is defined as follows according to (10) in SI References.

(a) the unique unit vector that lies in the plane containing $K_h^{e,-}$;

(b) $n_h^-(x) \cdot (x-y) \geq 0,\ \forall y \in K_h^{e,-} \cap B_\epsilon(x)$, where $B_\epsilon(x)$ is a ball centered in $x$. The radius $\epsilon(>0)$ of $B_\epsilon(x)$ is sufficiently small so that $\epsilon << |e|$, the length of edge $e$.

The conormal $n_h^+$ is defined similarly. With this definition, we have $n_h^+ \neq n_h^-$ in general. Let $P_k(D)$ denote the space of polynomials of degree not greater than $k$ on any planar domain $D$. The discrete DG space $S_{h,k}$ of scalar function associated with $\Gamma_h$ is $S_{h,k} = \left\{\chi \in L^2(\Gamma_h): \chi|_{K_h} \in P_k(K_h),\ \forall K_h \in T_h\right\}$, i.e., the space of piecewise polynomials which are globally in $L^2(\Gamma_h)$. The vector-valued DG space $\Sigma_{h,k}$ associated with $\Gamma_h$ is chosen to be $\Sigma_{h,k} = \left\{\varphi \in [L^2(\Gamma_h)]^3: \varphi|_{K_h} \in [P_k(K_h)]^3,\ \forall K_h \in T_h\right\}$. For $v_h \in S_{h,k}$ and $r_h \in \Sigma_{h,k}$, we use $v_h^\pm$ and $r_h^\pm$ to denote the trace of $v_h$ and $r_h$ on $e = K_h^{e,-} \cap K_h^{e,+}$ taken within the interior of $K_h^{e,-}$ and $K_h^{e,+}$, respectively. We refer readers to (13) for the definitions of the surface gradient operator $\nabla_\Gamma$ and the laplace-Beltrami operator $\Delta_\Gamma$ on $\Gamma$. By introducing the auxiliary variable $q = \sqrt{D}\nabla_\Gamma a$, the model problem Eqs. (1) can be rewritten as a first order system of equations:

$$a_t + \nabla_\Gamma \cdot \left(-\sqrt{D} q\right) = f(a)\,,$$
$$q - \nabla_\Gamma g(a) = 0\,,$$
$$\frac{db}{dt} = k_4(\bar{a} - k_{ss})b\,,$$
$$f(a) = \frac{k_0}{1+(\beta u)^{-q}} + \frac{k_1}{1+(\gamma p a)^{-h}} - k_2 a - k_3 b a\,,\ \bar{a} = \frac{\oint\ a ds}{\oint\ 1 ds},\ p = \frac{1}{1+(\beta u)^{-q}}\,.$$

where $g(a) = \sqrt{D}a$. The semi-discrete LDG for solving the above equations is defined by: Find $a_h \in S_{h,k}$ and $q_h \in \Sigma_{h,k}$, such that for all test functions $v_h \in S_{h,k}$ and $r_h \in \Sigma_{h,k}$,

$$\begin{aligned} &\int_{K_h} (a_h)_t v_h\, dx - \int_{K_h} \left(-\sqrt{D} q_h\right) \cdot \nabla_{\Gamma_h} v_h\, dx - \int_{\partial K_h} \widehat{\sqrt{D} q_h} \cdot n_h v_h\, dx = \int_{K_h} f v_h\, dx \\ &\int_{K_h} q_h \cdot r_h\, dx = -\int_{K_h} g(a_h) \nabla_{\Gamma_h} \cdot r_h\, dx + \int_{\partial K_h} \hat{g} n_h \cdot r_h\, dx\,, \qquad (S1) \\ &\frac{db_h}{dt} = k_4\left(\underline{a}_h - k_{ss}\right) b_h\,, \\ &\underline{a}_h = \frac{\oint_{\Gamma_h} a_h\, ds}{\oint_{\Gamma_h} 1\, ds}, \qquad p = \frac{1}{1+(\beta u)^{-q}}\,. \end{aligned}$$

Here $\hat{g}$ and $\widehat{\sqrt{D} q_h}$ are numerical fluxes which will be described later on. To facilitate definitions of numerical fluxes, following trace operators $\{\cdot\}$ and $[\![\cdot]\!]$ are introduced by following ideas in (10).

**Definition.** Denote by $K_h^{e,-}$ and $K_h^{e,+}$ the two elements sharing the common edge $e$. For $v \in L^2(\Gamma_h)$, $\{v\}$ and $[\![v]\!]$ are defined as $\{v\} = \frac{1}{2}(v^- + v^+)$, $[\![v]\!] = v^+ - v^-$ on $e$. For $\varphi \in [L^2(\Gamma_h)]^3$, $\{\varphi, n_h\}$ and $[\![\varphi, n_h]\!]$ are defined as $\{\varphi, n_h\} = \frac{1}{2}(\varphi^+ \cdot n_h^+ - \varphi^- \cdot n_h^-)$, $[\![\varphi, n_h]\!] = \varphi^+ \cdot n_h^+ + \varphi^- \cdot n_h^-$ on $e$.

Denote by $S_{K_h^+}^{K_h^-} \in \{0,1\}$ a switch function (14). $S_{K_h^+}^{K_h^-}$ is associated with $K_h^+$ on the edge that $K_h^+$ and $K_h^-$ share, and is defined by:

$$S_{K_h^+}^{K_h^-} = \begin{cases} 1\,, & if\ N_{K_h^+} > N_{K_h^-} \\ 0\,, & otherwise \end{cases}\,.$$

The numerical fluxes on the edge $e$ for $K_h^{e,-}$ and $K_h^{e,+}$ are defined respectively, as follows. The diffusive fluxes $\left(\widehat{\sqrt{D}q}^+, \hat{g}^+\right)^T$ and $\left(\widehat{\sqrt{D}q}^-, \hat{g}^-\right)^T$:

$$\widehat{\sqrt{D}q}^+ = \left(\frac{[\![g(a_h)]\!]}{[\![a_h]\!]}\{q_h, n_h\} - C_{11}[\![a_h]\!] + C_{12} \cdot n_h^+ [\![q_h, n_h]\!]\right) n_h^+,$$

$$\widehat{\sqrt{D}q}^- = -\left(\frac{[\![g(a_h)]\!]}{[\![a_h]\!]}\{q_h, n_h\} - C_{11}[\![a_h]\!] + C_{12} \cdot n_h^+ [\![q_h, n_h]\!]\right) n_h^-;$$

$$\hat{g}^+ = \{g(a_h)\} - C_{12} \cdot n_h^+ [\![a_h]\!],$$

$$\hat{g}^- = \{g(a_h)\} - C_{12} \cdot n_h^+ [\![a_h]\!].$$

Here the penalization coefficients $C_{11}$ and $C_{12}$ are chosen to be

$$C_{11} = \frac{1}{h_e}, \quad C_{12} = \frac{1}{2}\left(S_{K_h^{e,+}}^{K_h^{e,-}} n_h^+ + S_{K_h^{e,-}}^{K_h^{e,+}} n_h^-\right).$$

With the above choices of the numerical fluxes it yields that $[\![\hat{g}]\!] = 0$ and $[\![\widehat{\sqrt{D}q}, n]\!] = 0$. Thus these numerical fluxes are consistent and conservative. Moreover, they allow for a local resolution of $q_h$ in terms of $a_h$. The second-order accurate TVD Runge-Kutta (RK) time discretization is used to solve the semi-discrete scheme $(S1)$, which can be formulated as an ordinary differential equation:

$$\Phi_t = L(\Phi, t). \qquad (S2)$$

The second-order accurate TVD RK method for solving Eq. $(S2)$ is given by

$$\Phi^{(1)} = \Phi^n + \Delta t_n L(\Phi^n, t_n), \quad \Phi^{n+1} = \tfrac{1}{2}\Phi^n + \tfrac{1}{2}\Phi^{(1)} + \tfrac{1}{2}\Delta t_n L\left(\Phi^{(1)}, t_{n+1}\right).$$

Here $\Delta t_n$ is the time step size. In this paper, we only considered planar triangulations of the surface which are at most second-order accurate. Therefore, we choose the second-order accurate TVD RK time-stepping method. Also, the polynomial degree $k$ of the DG spaces used in this work is 1. Higher-order accurate surface approximation is needed in order to improve the overall accuracy of the scheme. We refer readers to (9) for accuracy tests of this numerical scheme.

**Movie Files:**

1. **Spherical bud**
2. **Tubular bud**

**SI References.**


1. M. Jin, *et al.*, Divergent Aging of Isogenic Yeast Cells Revealed through Single-Cell Phenotypic Dynamics. *Cell Syst* **8**, 242-253.e3 (2019).
2. Y. Li, *et al.*, A programmable fate decision landscape underlies single-cell aging in yeast. *Science* **369**, 325–329 (2020).
3. Z. Zhou, *et al.*, Engineering longevity—design of a synthetic gene oscillator to slow cellular aging. *Science* **380**, 376–381 (2023).
4. Y. Li, *et al.*, Multigenerational silencing dynamics control cell aging. *Proc Natl Acad Sci U A* **114**, 11253–11258 (2017).
5. C.-S. Chou, Q. Nie, T.-M. Yi, Modeling Robustness Tradeoffs in Yeast Cell Polarization Induced by Spatial Gradients. *PLoS ONE* **3**, e3103 (2008).
6. K. Tsai, *et al.*, Role of combined cell membrane and wall mechanical properties regulated by polarity signals in cell budding. *Phys Biol* **17**, 065011 (2020).
7. K. D. Moran, D. J. Lew, How Diffusion Impacts Cortical Protein Distribution in Yeasts. *Cells*

**9**, 1113 (2020).
8. E. Dague, *et al.*, An atomic force microscopy analysis of yeast mutants defective in cell wall architecture. *Yeast* **27**, 673–684 (2010).
9. S. Xu, Z. Xu, Local Discontinuous Galerkin Methods for Solving Convection-Diffusion and Cahn-Hilliard Equations on Surfaces (2024) https:/doi.org/10.48550/ARXIV.2401.02069 (February 16, 2024).
10. P. F. Antonietti, *et al.*, High Order Discontinuous Galerkin Methods for Elliptic Problems on Surfaces. *SIAM J. Numer. Anal.* **53**, 1145–1171 (2015).
11. B. Goldenbogen, *et al.*, Dynamics of cell wall elasticity pattern shapes the cell during yeast mating morphogenesis. *Open Biol.* **6**, 160136 (2016).
12. J. Schaber, *et al.*, Biophysical properties of Saccharomyces cerevisiae and their relationship with HOG pathway activation. *Eur. Biophys. J.* **39**, 1547–1556 (2010).
13. G. Dziuk, C. M. Elliott, Finite element methods for surface PDEs. *Acta Numer.* **22**, 289–396 (2013).
14. J. Peraire, P.-O. Persson, The Compact Discontinuous Galerkin (CDG) Method for Elliptic Problems. *SIAM J. Sci. Comput.* **30**, 1806–1824 (2008).
